# Cell heterogeneity contributes to the variable response of HIV-1 to latency reversing agents

**DOI:** 10.1101/2025.11.26.690455

**Authors:** Rachel Topno, Hussein Karaki, Flavia Mazzarda, Oriane Pourcelot, Manon Philippe, Kazem Zibara, Ovidiu Radulescu, Edouard Bertrand

## Abstract

Transcriptional noise contributes to gene expression variability, but its origin and impact in HIV-1 latency remain incompletely characterized. Here, we combine a dual-copy MS2-tagging system with novel mathematical analysis to investigate the variability of HIV-1 transcription in live cells. In the basal state, the transcriptional activity of proviruses located in the same cell was uncorrelated, indicating that expression variability primarily comes from the intrinsic stochasticity of promoter dynamics. Upon stimulation with diverse latency-reversing agents, viral transcription became more correlated within cells than across them, revealing a shift from promoter-driven to cell state–driven variability. Analysis of the molecular drug targets confirmed variable effects across cells. Our findings indicates that cellular heterogeneity shapes the response to latency reversing agents and demonstrate how quantitative tools can dissect noise sources. This work offers mechanistic insights into HIV-1 latency and informs strategies to target the latent viral reservoir.

**Teaser:** Live transcription imaging reveals how cell heterogeneity contributes to the variability of HIV-1 activation.

## Introduction

Live-cell imaging studies have shown that transcription does not proceed at a uniform rate even under steady-state conditions. Instead, it occurs in bursts, during which many mRNAs are rapidly transcribed, followed by periods of transcriptional silence (1). This burst-like behavior reflects the stochastic nature of promoter activity, with variable durations of active and inactive periods, an effect commonly referred to as transcriptional noise (reviewed in 1–5). Importantly, in a population of clonal cells exhibiting bursting behavior, this noise manifests as heterogeneity: some cells appear transcriptionally active, while others remain silent. This heterogeneity can have important phenotypic consequences (see 2–5 and therein). Understanding the various sources of transcriptional noise is a fundamental but yet incompletely resolved question (6).

Transcriptional noise has two major sources: extrinsic and intrinsic, which are usually distinguished by considering the behavior of identical copies of a gene within a cell (7–9). Intrinsic noise is mainly generated by the stochasticity of molecular events arising at a single transcription site, such as transcription factors binding-unbinding, nuclear architecture modifications (DNA looping, promoter-enhancer interaction, nuclear compartmentalization), local chromatin changes (nucleosome occupancy, histone modifications, occupancy of regulatory elements), polymerase pausing and DNA supercoiling (10,11). Intrinsic noise affects various gene copies within a cell in an independent, uncorrelated manner (12) and it is generally modeled as a promoter transitioning stochastically between transcriptionally active and inactive states (13). Single molecule studies such as single molecule DNA footprinting or electron microscopy were able to reveal and characterize these molecular states (14,15).

In contrast to intrinsic noise, extrinsic noise affects the copies of a gene within the same cell in a correlated manner. It can result from cell-to-cell fluctuations in the concentration of transcription factors or molecules involved in signal transduction, heterogeneity in the metabolic cellular status, cell-cycle stage, or in the cellular environment (16–19). Identifying and separating the two types of noise is crucial because extrinsic noise can introduce confounding effects if only intrinsic noise is considered (20). Clarifying the contributions of extrinsic and intrinsic sources of noise is also particularly important in biological systems where transcriptional noise has phenotypic consequences. This is the case of HIV-1 (19,21 for review), where stochastic variation of promoter activity has been proposed to control latency entry and exit, as well as the heterogeneous response of cells to latency reversing agents (LRAs), which limits current curing strategies (22,23).

The HIV-1 genome consists of a single transcription unit that is integrated into cellular chromatin. Transcription is carried out by the cellular machinery, resulting in primary transcripts that remain unspliced to be packaged into virions, or that are alternatively spliced to produce the various viral mRNAs and proteins. The HIV-1 promoter contains binding sites for a number of transcription factors including NF-κB, SP1 and TBP that binds the TATA box. NF-κB activates viral transcription in response to extracellular signals such as the TNFα cytokine, while SP1 is required for the basal and activated transcriptional activity of the virus (24–28). The virus also encodes the Tat protein, which stimulates viral transcription by more than 100 fold and triggers a positive feedback loop. Tat binds to the nascent TAR RNA together with P-TEFb, which enables the release of RNA polymerases from a promoter proximal pause (29,30). Critically, the HIV-1 virus can exist in a latent and an acute state, which are mainly controlled at the transcriptional level. In latent cells, viral transcription is minimal and the Tat feedback loop is OFF. In acutely infected cells, viral transcription is active, producing high levels of Tat and fully activating the viral promoter. The discovery of transcriptional noise has generated considerable interest in HIV-1 biology, particularly in relation to its role in latency entry and exit. Early studies using GFP-tagged mini-viruses suggested that these viruses behave as bi-stable switches, which stochastically alternate between meta-stable ON and OFF states resembling acute and latent states, respectively (31–35). Direct imaging of HIV-1 transcription using MS2-tagged reporters in living cells further showed that the viral promoter stochastically switches between active and inactive states as a result of gene bursting (36,37). In absence of Tat, this bursting could potentially trigger latency exit by inducing the Tat feedback loop.

In patients, HIV infection can be controlled with highly active antiretroviral therapy (HAART). However, interruption of treatment leads to viral rebound within weeks and lifelong treatment is required, posing risk of secondary drug effects and the emergence of resistant strains (38). The viral rebounds are primarily driven by latently infected cells, which form the viral reservoir. This reservoir is established early in infection and may be maintained by low-level viral propagation, even under antiretroviral therapy (39). Importantly, the half-life of the reservoir is estimated to be around 44 months, meaning its eradication could require more than 60 years of continuous treatment (22,23,38). This highlights the need for new strategies that target the reservoir for the development of an effective cure. One approach is the "shock and kill" strategy (40) whose goal is to reactivate viral transcription using latency-reversing agents (LRAs), thereby allowing the immune system to recognize and eliminate latent cells. Unfortunately, this approach is currently limited by the inability to activate all latent viruses, thus hindering their complete eradication (see (41) and therein). This inability is due to the stochastic response of latent viruses to LRAs, with some viruses activating transcription and others not, even in clonal populations (42,43). Importantly, this variability has been observed in model systems and also directly demonstrated in patient cells (21,44).

While the stochastic viral responses are a key limitation for therapies targeting the reservoir, we only have a partial understanding of their origins. In particular, it is unclear if they stem from intrinsic or extrinsic sources. The intrinsic stochastic dynamics of latent promoters would cause them to dynamically switch between different molecular states, some of which are capable of reactivation and others that are not. Similarly, heterogeneities between cells would also generate variability in how individual cells respond to LRAs. More generally, understanding the interplay between noise sources and latency would provide valuable insights into the factors that influence reservoir dynamics in patients.

Here, we developed a system to image in real time the transcription of two identical viral copies integrated into human cells. We quantitatively dissected the respective contributions of intrinsic and extrinsic sources of transcriptional heterogeneity under basal conditions and upon stimulation with diverse LRA treatments. We show that the heterogeneity in basal viral transcription is entirely driven by intrinsic noise, related to promoters being in different molecular states, while the heterogeneity in the response to LRAs shows a large contribution of extrinsic factors related to the variable properties of individual cells. Cellular heterogeneity is thus an important issue for therapeutic approaches targeting the viral reservoir, and likely plays an important role in the reservoir dynamics in patients.

## Results

### Generation of a two-copy HIV-1 system to measure the contribution of extrinsic and intrinsic sources of noise

Inserting multiple copies of a reporter gene into the same cell and measuring the correlation of their expression provides a robust way to quantify the relative contributions of intrinsic and extrinsic noise (9). To construct such a system, we generated two identical HIV-1 vectors bearing different selection markers and introduced them at a precise location in the human genome by CRISPR/Cas9 mediated homologous recombination. We chose an integration site in the TBC1D5 gene, which was previously identified as a viral integration site in a patient (45), and the viral vector that we integrated was similar to the one we used previously to image HIV-1 transcription (36,37; Figure 1A). This vector harbored the 5’ LTR with the viral promoter and TAR element, the splicing donor SD1, splice acceptor A7 and the 3’ LTR with the 3’-end processing and polyadenylation site. We used the MS2/MCP system to image viral RNA in living cells (46) and inserted 128 MS2 loops in the intron between SD1 and A7 (36). We previously showed that this intron splices post-transcriptionally and enables imaging transcription independently from splicing (36). The 128xMS2 tag also enables the visualization of single RNAs for extended time periods and we used the newly developed MCP-tdStayGold variant to further improve RNA imaging (47). Overall, this system allows visualization of single RNAs with exceptional photostability and brightness, ensuring the detection of every transcriptional event over long time periods. Importantly, the viral vector did not express Tat and thus corresponded to a transcriptional status analogous to the one of latent cells.

**Figure 1.**
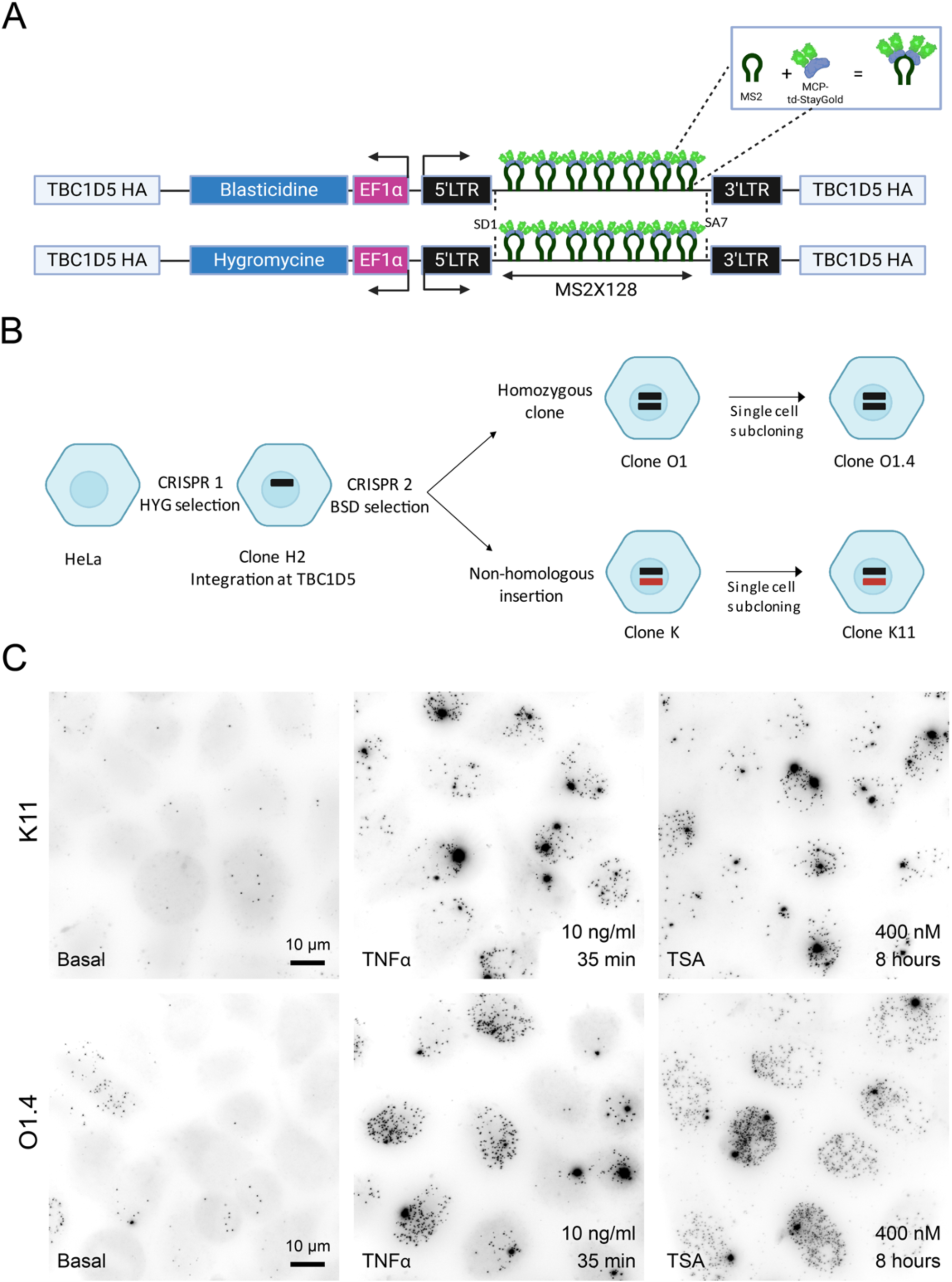
Generation and characterization of two-copy HIV-1 CRISPR clones. **A.** Schematic representation of two CRISPR repair cassettes designed to insert an HIV-1 reporter containing 128 MS2 stem-loops in the TBC1D5 gene. Key components of the reporter include the major HIV-1 splice donor site (SD1), the last HIV-1 splice acceptor site (SA7), homology arms targeting TBC1D5 (TBC1D5 HA), and the selection markers blasticidin and hygromycin. The human elongation factor 1 alpha (EF1α) promoter drives expression of the selectable marker. The MS2 coat protein (MCP-tdStayGold; green) binds to the MS2 stem-loops. **B**. Schematic outlining the experimental workflow for generating the CRISPR-edited HeLa clones. Two rounds of CRISPR transfections and clone selection were performed. In the first round, a cassette with the hygromycin selection marker was used, resulting in the H2 clone. A second round of CRISPR transfection using the blasticidine marker was performed on the H2 clone, yielding two subclones: the O1 homozygous clone and the K clone with a non-homologous integration site. Both clones were subsequently sub-cloned to produce the O1.4 and K11 clones, respectively. **C**. The micrographs represent smiFISH microscopy images of clone K11 (top panel) and O1.4 (bottom panel) under Basal, TNFα and TSA stimulated conditions (from left to right). The concentration and duration of treatment are indicated in the bottom-right corner of each image.

The two copies of the MS2-tagged viral reporter were inserted in the genome sequentially using two rounds of CRISPR editing in HeLa cells (Figure 1B). In the first round, we used the repair cassette bearing the hygromycin selectable maker, and after PCR genotyping different clones, we identified a correct integration in clone H2 (Figure S1B). After subcloning, this cell line was further modified by CRISPR with the second repair cassette carrying the blasticidin selection maker, and two clones were characterized in detail (Figure 1B and S1B). The first clone, referred to as O1.4, carried a correct integration for the second CRISPR and was thus homozygous with the two viral reporters integrated in the two alleles of the TBC1D5 gene. The other clone, K11, had the second repair cassette integrated in a different site, and was thus heterozygous with respect to the TBC1D5 gene. The K11 clone allowed us to evaluate the contribution of extrinsic noise independently of the integration site.

The two clones were subjected to single cell sorting and subcloned to ensure homogeneity, and analyzed by fluorescent microscopy to assess the quality of RNA imaging. Viral transcription was activated and RNA expression was assessed with smiFISH using probes hybridizing to the MS2 sequences. We observed two bright spots corresponding to the two expected viral transcription sites, as well as many dimmer spots corresponding to single RNAs (Figure S1A). Both colocalized with signals from the MCP-tdStayGold, indicating the accuracy and sensitivity of the RNA imaging system (Figure S1A, Supplementary Videos).

### SmFISH indicates a lack of correlation between HIV-1 copies in the basal state and moderate correlation during promoter activation

To determine the transcriptional activity of the two integrated copies of the MS2-tagged reporter, we performed smiFISH under basal and activated conditions and quantified the resulting signals. Specifically, we utilized two well-known transcriptional inducers of HIV-1, tumor necrosis factor-alpha (TNFα) and Trichostatin A (TSA, Figure 1C). TNFα activates NF-kB, which directly binds the HIV-1 promoter (48–50), while TSA inhibits histone deacetylases and promotes the formation of an open chromatin (51). Under basal conditions, smiFISH revealed low levels of nascent RNAs from either allele, in both clones K11 and O1.4 (0.3 and 0.4 RNA per transcription site; Figure 2), yielding an average of 2.7 and 4.2 molecules of released RNAs per cell, respectively (Figure S2). Following treatment with TNFα for 35 minutes and TSA for 8h, we observed a marked increase in transcriptional output, as evidenced by a higher number of nascent and released RNAs for both the K11 and O1.4 clones, and for both reporter copies in each clone (Figure 2 and S2). TNFα treatment resulted in a 25-fold increase in nascent RNAs for K11 and 22 fold in O1.4, and a 7 fold increase in released RNAs in K11 and 6 fold in case of O1.4. TSA led to a 36 and 12 fold increase in nascent RNA for K11 and O1.4, respectively, and a 13 and 14 fold increase in released RNAs. This was in line with the expected response to these transcriptional inducers (27,52–54). These results demonstrate that the CRISPR-integrated HIV-1 system is functionally responsive to known transcriptional inducers, validating the system’s utility for monitoring viral transcription in live cells.

**Figure 2.**
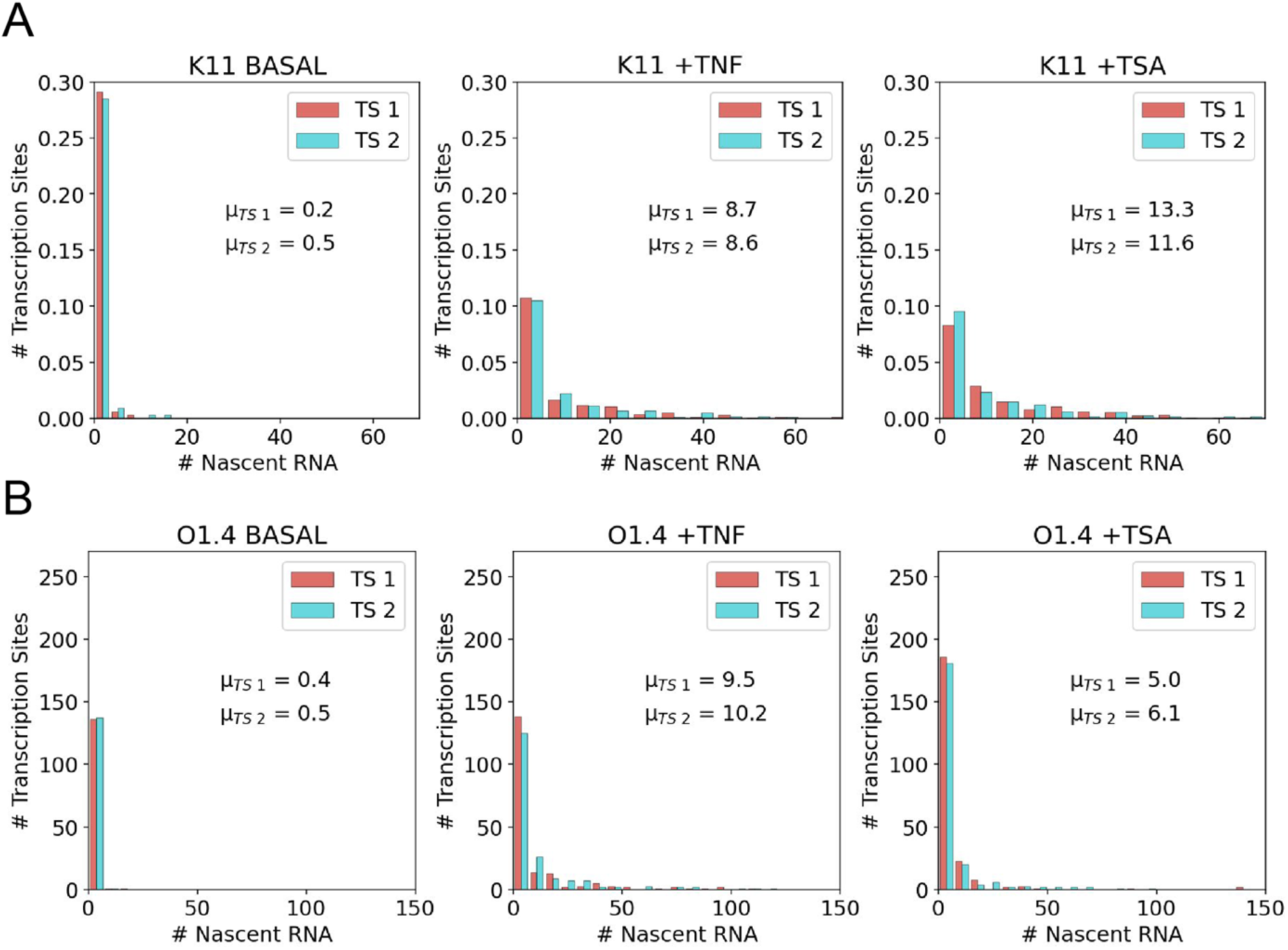
Transcriptional activity of the two HIV-1 copies under basal condition and during activation. **A-B**. The graphs display the distribution of nascent RNA counts per transcription site (TS) for the two reporter copies, distinguished by color (red and blue), for K11 (A) and O1.4 (B). The mean nascent RNA count for each transcription site (denoted as µTS1 and µTS2) is provided in the center of the plot. The x-axis represents the number of nascent RNA counts, while the y-axis shows the number of transcription sites exhibiting those counts. The number of cells in clone K11 used for Basal, TNFα and TSA treatments are 99, 301, and 228 respectively, while the number of cells used for Basal, TNFα and TSA treatments are 138, 188, and 226 respectively in clone O1.4.

Next, we compared the activities of the two HIV-1 copies located in the same cell. Scatter plots comparing the number of nascent RNAs for the two copies showed an absence of correlation in the basal state, for both K11 and O1.4 (Figure 3A and 3B). In contrast, a weak correlation was observed in the K11 clone 35 minutes after TNFα addition and a higher one was seen 8h after TSA addition (Pearson correlation coefficient of 0.15 and 0.26, respectively). For the O1.4 clone, no correlation was seen after TNFα addition and a weak one after TSA treatment. We then measured the total, extrinsic and intrinsic noise using the total law of variance (9). We found that the total noise levels decreased after transcriptional induction, for both K11 and O1.4 clones (Figure 3C). Meanwhile, the proportion of extrinsic noise increased, reaching up to ∼35% in clone K11 after TSA addition (Figure 3D), and greater than 20% for the O1.4 clone (Figure 3E).

**Figure 3.**
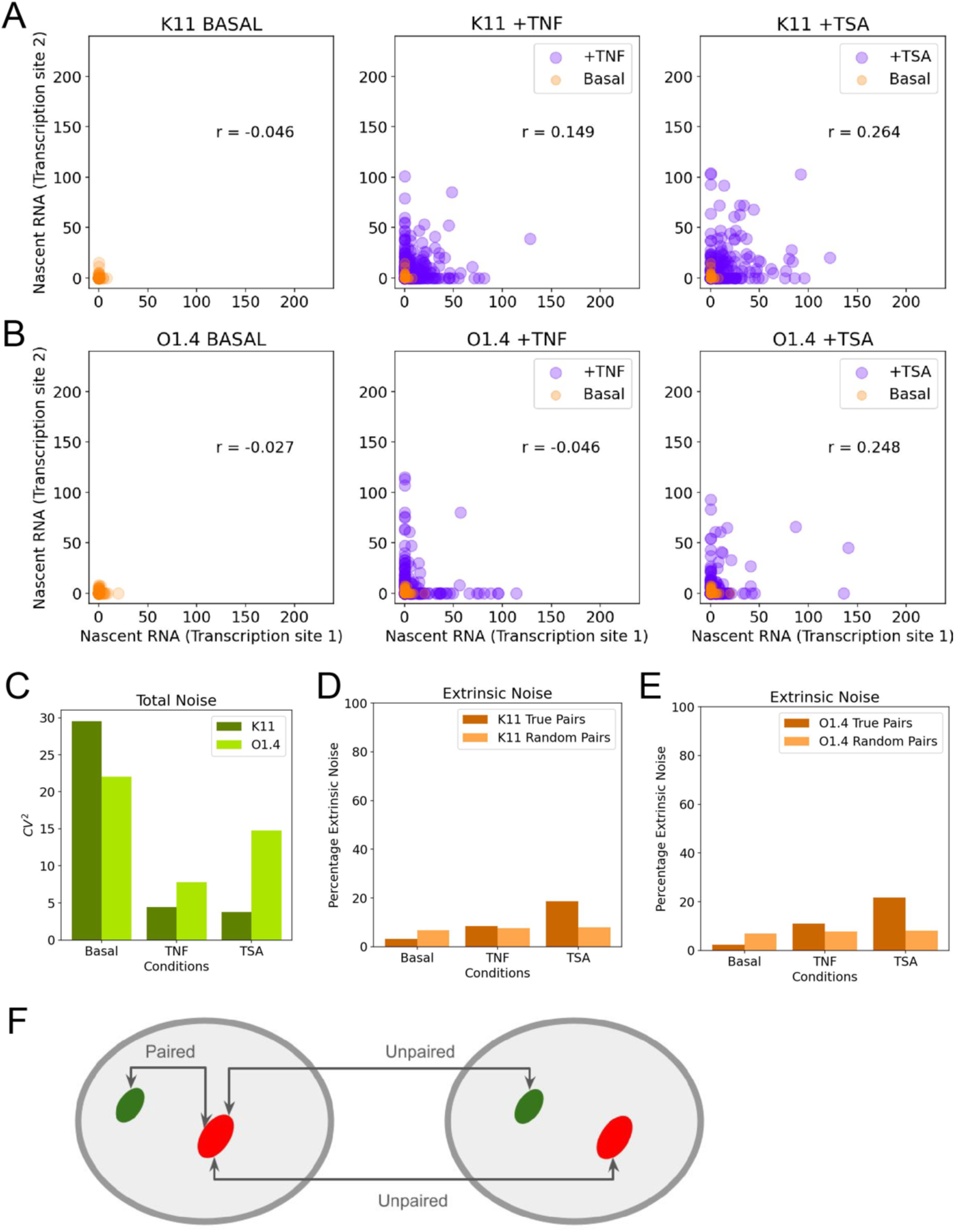
SmiFISH analysis reveals the presence of extrinsic noise and cellular heterogeneity. **A-B.** Correlation scatter plots for nascent RNA counts from transcription site 1 (x-axis) and transcription site 2 (y-axis) for clones K11 and O1.4, respectively. The Pearson correlation coefficient (r) is displayed in the center of each plot. From left to right, the plots represent the basal, TNFα, and TSA treatment conditions. **C**. Bar plot showing the noise estimates (CV^2^) for clones K11 and O1.4 across various conditions (basal, TNFα, TSA) on the x-axis. The colors representing each clone are indicated in the legend located at the top right of the panel. **D-E**. Bar plot depicting the percentage of extrinsic noise for clones K11 and O1.4 under basal, TNFα, and TSA treatment conditions. The colors corresponding to each condition are provided in the legend at the top right of the panel. **F.** Schematic representation depicting the paired and unpaired transcription sites.

The extrinsic noise could originate from variations in the extracellular environment or from heterogeneities in the ability of single cells to respond to the inducer. To discriminate between these two possibilities, we designed a procedure to quantify separately these sources of noise (Figure 3F). We applied the total variance decomposition to two types of reporter copies: those placed in the same cell (paired transcription sites, referred to as ‘cellular’ extrinsic noise; Figure 3D and 3F) and those placed in different cells (unpaired transcription sites, referred to as ‘environmental’ extrinsic noise; Figure 3E and 3F). For both clones, extrinsic noise of unpaired alleles was significantly lower after TNFα or TSA, as compared to paired alleles (Figure 3D and 3E), suggesting that a sizeable part of extrinsic noise following promoter activation was attributable to cell-to-cell heterogeneity. Overall, these data demonstrate that both copies of the integrated MS2 reporter were transcriptionally active and responsive to external stimuli, and that extrinsic noise was readily detectable upon transcriptional stimulation, with a component arising from cellular heterogeneity.

### Live cell imaging reveals a rapid transient transcriptional stimulation after TNF**α**, and a slow, continuous increase after TSA treatment

To characterize in detail the transcriptional fluctuations of the HIV-1 promoter, we imaged the two-copy cell lines live. We recorded time-lapse movies for 15 hours at a rate of one 3D image per minute, using an illumination power enabling the detection of single RNAs throughout the movies (Figure 4A and 4B, Supplementary Videos). In both basal and after drug treatments, cells exhibited varying and fluctuating levels of transcriptional activity, as illustrated in the snapshots extracted from the movies (Figure 4A and 4B). To further analyze the viral transcriptional behavior, raw live-cell imaging data were processed to extract transcription site intensities (Figure 4C). Single RNAs present in the nucleoplasm were used to calibrate the signals such that the exact number of nascent RNAs was calculated at each time point. Examples of trajectories from single cells in the basal condition showed a typical bursting pattern of transcriptional activity, with little apparent temporal correlation between the bursts of the two reporter copies, in either K11 or O1.4 clones (Figure 4D and 4E). A higher transcriptional activity was visible following stimulation with TNFα, TSA and JQ1, while still highly heterogeneous over time and between cells (Figure 4D and 4E). Heatmaps displaying the activities of many cells and their averages showed that transcriptional activation raised rapidly after TNFα treatment and culminated after about 30 minutes with a maximal activation of 16 fold, before returning to basal levels after an additional 30 minutes (Figure 5A and 5B). In contrast, TSA led to a gradual and sustained increase in transcriptional activity over time, which culminated at about 10h for clone O1.4 and 15h for clone K11, likely reflecting clonal differences in TSA sensitivity. The maximal fold activation was 19 and 7 fold for clone K11 and O1.4, respectively, in line with the responses observed by smiFISH. We also tested JQ1 and PMA-Ionomycin, two other traditional activators of HIV-1. JQ1 is an inhibitor of BET proteins such as BRD4. It blocks their binding to acetylated lysines and has been shown to be a potent activator of latent HIV-1 viruses (55). PMA-Ionomycin activates protein kinase C and increases calcium levels, which activates viral transcription via NF-kB, NFAT and AP1 (56,57). Upon addition of JQ1 in K11 cells, a rapid and sustained increase in transcriptional activity was observed, starting approximately 30 minutes after addition of the drug, reaching a peak at 95 minutes and then plateauing for the rest of the movie. In contrast, treatment with PMA-Ionomycin resulted in weak activation with RNA levels remaining close to basal conditions. These observations highlight the diversity of transcriptional responses of HIV1-promoters, shaped by both the nature of the stimulus and, to a lesser extent, clonal variability.

**Figure 4.**
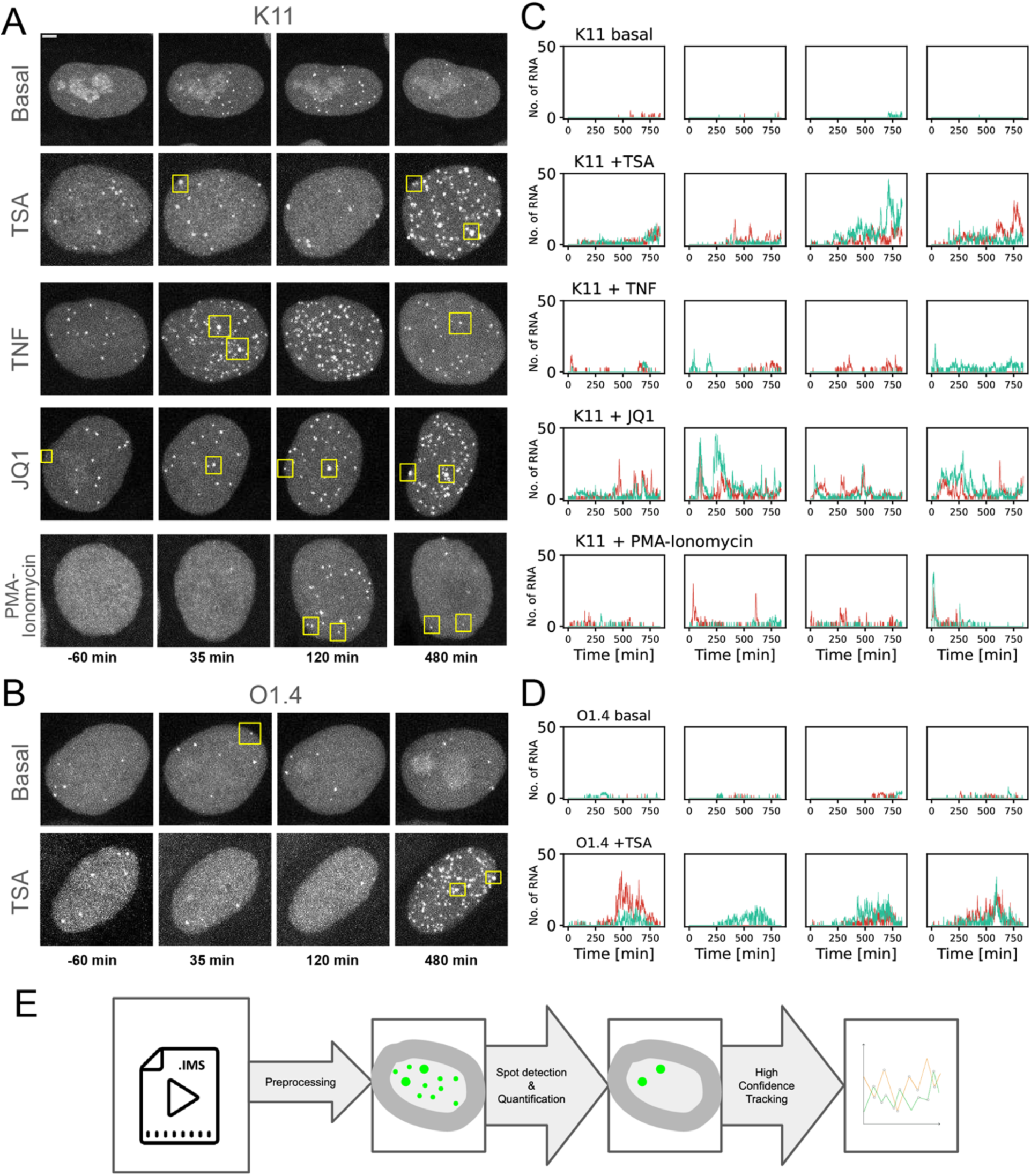
Live cell analysis of the response of the HIV-1 promoter to diverse activators. **A-B**. Snapshots from live imaging data showing a single nucleus at different timepoints for clones K11 and O1.4, respectively. From top to bottom, the conditions are Basal, TSA, TNFα, JQ1 and PMA-Ionomycin. Yellow squares highlight the activated transcription sites, visible as bright spots with intensities higher than the surrounding spots, where the latter are released single RNAs. Scale bar: 3 microns. **C-D**. Single cell trajectories of nascent RNA counts per transcription site over time for clones K11 and O1.4, respectively. For each clone, from top to bottom, the conditions are Basal, TSA, TNFα, JQ1 and PMA-Ionomycin treatment. The x-axis represents time (in minutes) over a total of 900 minutes (15 hours), and the y-axis shows the number of nascent RNA molecules. The two transcription sites corresponding to the reporter genes are depicted in red and green. **E**. Workflow illustrating the image analysis method for obtaining transcription site trajectories. The raw data undergoes preprocessing to obtain cropped movies for each nucleus, followed by spot detection, quantification, and high-confidence tracking. The final output is the transcription site RNA count over time.

**Figure 5.**
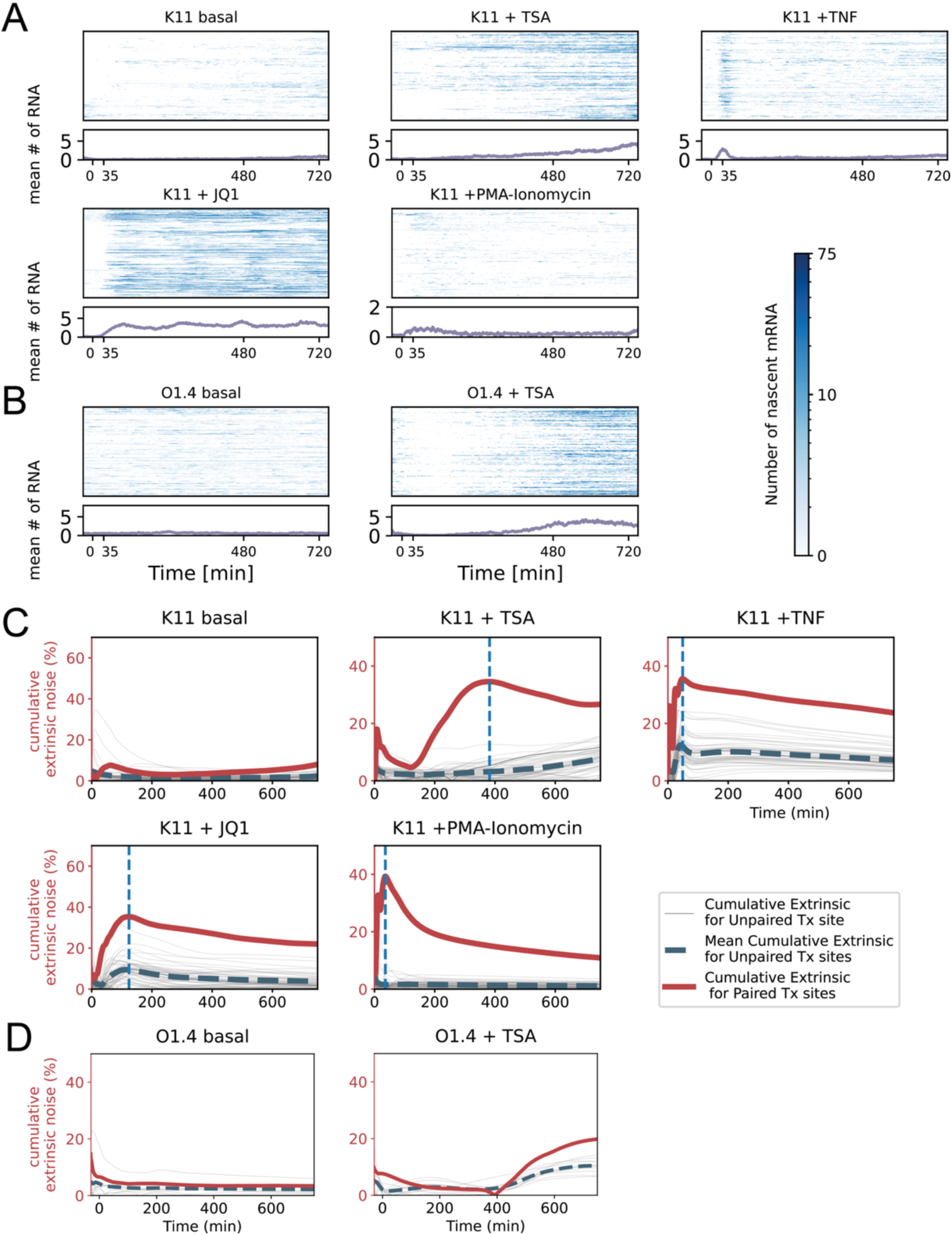
Analysis of drug-induced transcription reveals extrinsic noise and cell heterogeneity. **A-B**. Heatmaps displaying the nascent RNA count over time for clone K11 (A) and clone O1.4 (B), under Basal, TSA, TNFα, JQ1 and PMA-Ionomycin treatment conditions (as indicated). On the y-axis, each row represents the RNA count for an individual transcription site, with transcription sites within the same nucleus arranged in adjacent rows. The color bar maps the intensity of the color to the RNA count. Below each heatmap, the mean transcription site intensity is plotted over time. The number of cells analyzed was 39, 107, 76, 164, 142 for K11 in Basal, TNFα, TSA, PMA and JQ1, respectively, and 160 and 87 for O1.4 in Basal and TSA condition. **C-D**. Cumulative extrinsic noise as a percentage of total noise plotted over time for clones K11 and O1.4 under Basal, TSA, TNFα, JQ1 and PMA-Ionomycin treatment conditions. The bold red line represents the cumulative extrinsic noise for paired transcription sites as a percentage, while the dashed blue line denotes the averaged cumulative extrinsic noise for unpaired transcription sites. The thin grey lines represent individual cumulative extrinsic noise calculated for a number of randomly selected subpopulations of unpaired transcription sites. The vertical dashed blue line identifies the time where extrinsic noise is maximal in each condition.

### Cumulative analysis of transcription site activity over time reveals the extrinsic noise in response to promoter activation

Transcription site intensities at a given time provide only a snapshot of promoter activity, as nascent RNAs remain there for only a few minutes (36). In the non-stationary regime following application of the treatment, if some sources of noise have short time scales while others operate on long ones, it may be difficult to detect the latter using only transcription site intensities. To circumvent this issue, we integrated the signal of transcription sites over time and replaced the instantaneous signal with its corresponding cumulative area under the curve (cAUC). This quantity was used for computing the ‘cumulative’ extrinsic and intrinsic noise by applying the law of total variance to the time-integrated signal (see Methods, Figure 5C, 5D and S3). We also computed the ‘instantaneous’ values of noise by calculating noise values using the intensities at each time point (Figure S4A and S4B). Both the instantaneous and cumulative extrinsic noise was found to be low in basal condition for the K11 and O1.4 clone. It accounted for less than 5% of the total noise even after integrating the transcription site signals for 15h (Figure 5C and 5D). Thus, the intrinsic fluctuations of promoter activity were almost entirely responsible for expression heterogeneities in the basal state, at short and long timescales. In contrast, we observed a large increase in the cumulative extrinsic noise following TNFα treatment in the K11 clone, peaking at 36% of total noise after 35 minutes (Figure 5C). A similar result was obtained with PMA-Ionomycin, which also activates viral transcription via signaling pathways. Following the peak, cumulative extrinsic noise then slowly decreased, most likely because the RNAs produced shortly after drug addition provided most of the activity of the transcription sites even after much longer times. Interestingly, when the cumulative extrinsic noise was computed while starting the cumulative sum at a time point after the TNFα induced peak, it still increased, albeit slowly, reaching 12% after 10h (Figure S3). This indicated that besides the ∼1h peak of transcription activation, TNFα also has long term consequences on extrinsic noise. The initial increase in extrinsic noise seen immediately after TNFα addition was also visible in the instantaneous noise measurements although the data was much noisier (Figure S4A). JQ1 and TSA target chromatin and they led to similar effects as TNFα, but with slower dynamics. JQ1 showed a delayed peak in extrinsic noise at 120 min, with a maximal value of 35% of total noise (Figure 5C). The delay was much larger for TSA, which led to a late increase in the cumulative extrinsic noise (Figure 5C and 5D), peaking at 35% and 21% of total noise for clones K11 and O1.4, at 6h and 14h, respectively. Long-term effects of TSA addition are evident as a steady increase in cumulative extrinsic noise, even when the calculation began 9h after TSA was added (Figure S3). These results were confirmed by the instantaneous measures although these data were again much noisier (Figure S4).

As with fixed-cell images, we decomposed extrinsic noise into cellular and environmental components by computing the correlation of paired and unpaired transcription sites (Figure 5). In the basal state, the cumulative extrinsic noise was low for both paired and unpaired transcription sites, and for both the K11 and O1.4 clones (Figure 5C and 5D). Remarkably however, following stimulation with either any of the activators, and for both the K11 and O1.4 clones, the cumulative extrinsic noise for unpaired transcription sites (dashed blue plot) was several folds lower than for paired transcription sites (Figure 5C and 5D), indicating a low environmental extrinsic component compared to the cellular extrinsic one. The strongest differences were seen for K11, with 36% and 3% of total noise being extrinsic for paired and unpaired copies, respectively, at the peak of TSA treatment (400 minutes), 36% versus 12% after TNFα addition for 35 minutes (Figure 5C), 35% versus 9% for JQ1 and 40% versus 3% for PMA, respectively (Figure 5C). The effect was smaller for the O1.4 clone, but still important, with 25% versus 12% of noise being extrinsic for paired versus unpaired copies at the peak after TSA addition (Figure 5D). In summary, intrinsic noise dominates in the basal state, while drug treatments increase extrinsic noise with distinct dynamics for different signals. The higher noise in paired versus unpaired alleles indicates that these effects are mainly due to heterogeneity in the ability of single cells to respond to activators.

The analyses above were conducted on populations of cells. To better evaluate the activities of the two transcription sites in individual cells under different conditions, we plotted the cumulative intensities (cAUC) of the transcription sites belonging to the same cell against one another (Figure S5). A linear relationship was observed between the cumulative intensities of the two transcription sites, which is expected since both signals increase progressively over time. Interestingly, the slope was sometimes away from one. This suggests that the two reporter copies could display differential transcriptional capacity that could be sustained for up to 15h.

### Cross-correlation analysis extracts the time-scales of extrinsic noise

To better characterize the time-scale of the different noise components, we performed auto-correlation and cross-correlation analyses (See Materials and Methods). We computed the auto-correlation function of the signal of transcription sites, as well as the cross-correlation function between signals from pairs of sites within the same cell and across different cells. The decomposition laws also apply to these functions, enabling us to distinguish the auto-correlation of intrinsic and extrinsic noise. We further decomposed the extrinsic component into a cellular component and an environmental component using paired and unpaired transcription sites.

In the basal state, a minimal extrinsic component was confirmed for clones K11 and O1.4, as indicated by the alignment of paired and unpaired cross-correlation functions near zero (Figure 6A and B, left-most plots). The cross-correlation curves of paired reporters were flat in the basal conditions, indicating that promoter bursting rates were constant, as expected.

**Figure 6.**
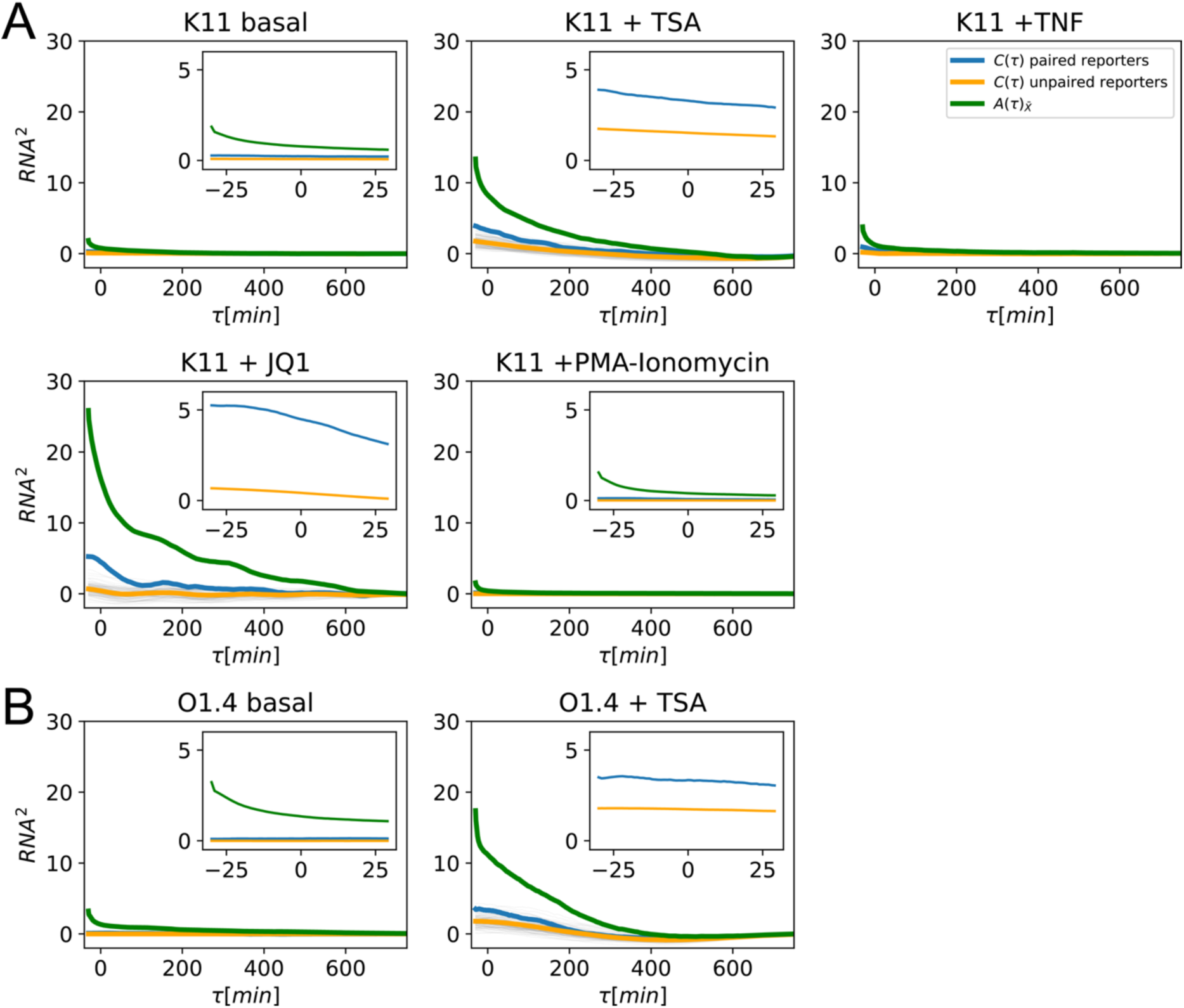
Cross-correlation analyses uncover the timescales of extrinsic and intrinsic noise. **A-B**. Auto- and cross-correlation functions for clones K11 and O1.4 across Basal, TSA, TNFα, JQ1 and PMA-Ionomycin treatments. The bold green line represents the auto-correlation function of the mean trajectories from two transcription sites within the same nucleus. The bold blue line denotes the cross-correlation of reporter pairs from the same nucleus, while the bold orange line represents the averaged cross-correlation of random reporter pairs sampled over several subpopulations of nuclei. Thin grey lines represent the cross-correlation for each such subpopulation of unpaired sites.

In treated cells, there was a difference between the paired (blue) and unpaired (orange) cross-correlation functions, indicating heterogeneities between cells in both clones and for all signals (Figure 6A and B). However, the establishment of cellular heterogeneity occurred on different timescales depending on the signal. To estimate these timescales, we used multi-exponential fitting of the difference between paired and unpaired cross-correlation curves. The fastest response was observed for PMA, with a single exponential fit yielding a timescale of 83 minutes. The next fastest response was for TNFα treatment, but in this case we found two exponentials corresponding to 93 and 200 minutes, respectively. JQ1 treatment induces heterogeneity on a unique long timescale of 195 minutes. The slowest response was for TSA, corresponding to 230 minutes in O1.4 and 352 minutes in K11 clones. This hierarchy of timescales was consistent with what we have obtained by the cumulative method in the previous analysis.

In summary, auto- and cross-correlation analyses revealed distinct timescales for extrinsic noise components, with minimal cellular variability in the basal state and drug-specific dynamics of cellular heterogeneity upon stimulation. TSA induced slow, long-term changes in promoter kinetics and cellular heterogeneity, while PMA-Ionomycin, TNFα and JQ1 triggered faster, more transient modulations, reflecting their differing regulatory effects on cells and the transcriptional machinery.

### Molecular analysis of drug targets reveal that drug treatment induces cell heterogeneities

Treating the K11 and O1.4 cells with TSA, TNFα, JQ1 and PMA-Ionomycin clearly show cell-to-cell differences in viral expression (Figure 2 and 5) and the appearance of cell heterogeneity, because viral copies in the same cell showed a higher correlation than copies in different cells (Figure 5C and Figure 6A). It is well known that TNFα acts through a complex signaling cascade activating the canonical NF-kB pathway (58–60). TSA, instead, modulates chromatin structure by inhibiting histone deacetylases I and II (HDACs) with a resulting increase of acetylated lysines (AcK) and active transcription (61,62). To gain some insights into the possible origin of the factors conferring heterogeneities in cellular properties, we analyzed the levels of several key components of these pathways in single cells, using immuno-fluorescence (IF) and FACS. As shown in Figure 7A, after 30 minutes treatment with TNFα, the NK-κB p65 protein translocated into the nuclei as expected. Interestingly however, the levels of nuclear p65 varied from cell to cell, indicating variable levels and/or timing in the activation of the pathway. Part of this may stem from the TNF receptor itself (TNFR1), as IF and FACS analysis showed that it was expressed to variable levels in cells at the start of the experiment (two folds difference between the third and first quartile). When analyzing the TSA pathway, lysine acetylation levels increased significantly in the nucleus after TSA treatment, with a regular increase from 2 to 8h of treatment (Fig. 7A and 7C). Interestingly, the cell-to-cell variability also dramatically increased, with a cell-to-cell standard deviation rising from ∼1 to ∼5 over 8h. The variability increased even after normalization to the mean, with a peak after 2h of TSA treatment (Figure 7E). The levels of acetylated lysine were also highly uneven with disproportionate representation of high intensity subpopulations, as shown by skewness values ranging from 1.81 to 2.01 and Gini index 0.45 to 0.62.

**Figure 7.**
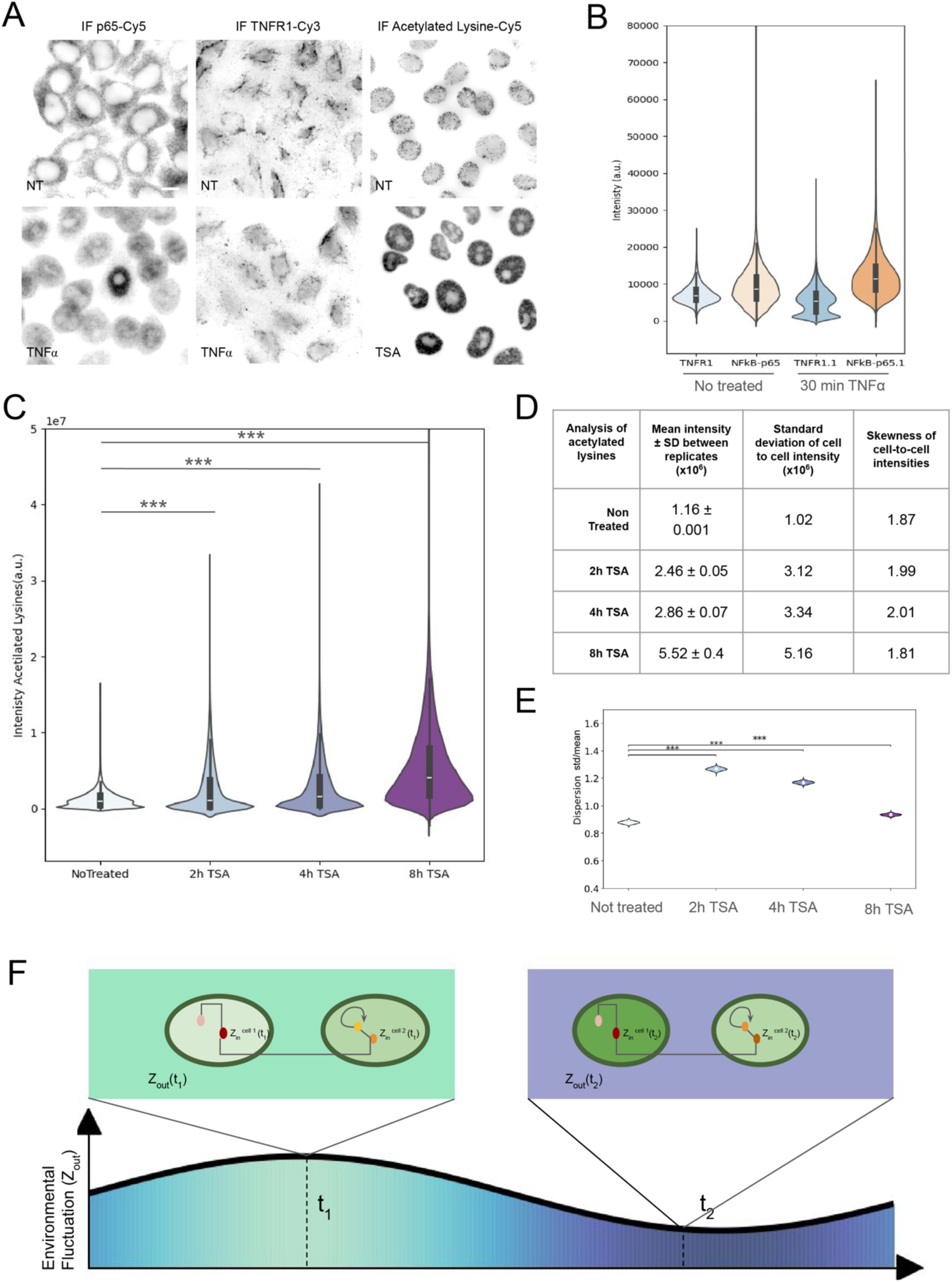
Molecular analysis of cellular heterogeneity. **A.** Representative immunofluorescence images of clone K11 under untreated conditions (upper panel) or following treatment with TNFα for 35 minutes or TSA for 8 hours (lower panel). From left to right, images show immunofluorescence staining (in grey) with antibodies against NF-κB p65, TNFR1, and acetylated lysines. Image contrast was adjusted individually to enhance visibility. Scale bar: 10 μm. **B.** Violin plot of cytometry data of K11 cells, untreated or treated for 30 minutes with TNFα and stained with TNFR1 or NFkB-p65 antibodies. The y-axes represent the fluorescence intensity measured for each cell (11496 cells for each sample). **C.** Violin plot of cytometry data of K11 cells, untreated or treated with TSA for the indicated time and stained with an antibody against acetylated lysines. The y-axis represents the intensity for each cell in two experiments (7913 cells for each experiment; Mann–Whitney U test: ***p<0.001). **D.** Recapitulative values of (i) mean intensity +/- the standard deviation (SD) calculated between replicates (n=2), (ii) standard deviation of cell-to-cell intensities and (iii) skewness of cell-to-cell intensities. values are calculated from data in C. **E.** Dispersion (standard deviation/mean) of the single cell intensities of acetylated lysines. The p-values are calculated using the two replicates and a bootstrapping algorithm (7000 samples per group without replacement, *** p<0.001). **F.** Schematic diagram of the different sources of extrinsic noise. The shaded plot represents the fluctuation in environmental extrinsic noise (Z_env_) shared by several cells, represented by the color of the plot and the color surrounding the cells. Two representative cells, X and Y, are shown at two distinct time points (t_1_ and t_2_) to depict cellular extrinsic noise 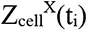 and 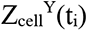, respectively, represented by the color inside the cells). These values reflect the distinct cellular conditions that two unpaired transcription sites may experience. Paired transcription sites share the same cellular and environmental conditions. Unpaired transcription sites experience the same environmental parameters but different intracellular conditions.

Altogether, these data show that cells display significant heterogeneity in key components of the NF-κB pathway. More remarkably, TSA treatment appears to generate heterogeneity within the cell population. These findings highlight that individual cellular properties contribute substantially to the observed variability in transcriptional activation and are themselves modified by drug treatment.

## Discussion

### Intrinsic promoter dynamics generates expression variability in the basal state

Several studies have used two-copy protein reporters in bacteria, yeast and mammalian cells (7,13,63–65). However, the use of protein reporters provides only an indirect measure of transcription, as the long half-lives of the mRNA and their encoded proteins (e.g., 2 hours for destabilized GFPs) tend to buffer transcriptional fluctuations. Real-time imaging of nascent transcripts, though currently a low-throughput method requiring extended observation time at high frame rate (>10h with one 3D image per minute), offers a direct measurement of extrinsic and intrinsic noise sources affecting transcription when combined with a two-copy reporter system (66). Here, we used cell lines carrying two copies of an HIV-1 reporter tagged with MS2 sites to measure their transcriptional activity in real time, in absence of the viral activator Tat. We found that basal transcription showed a bursting type of expression that was entirely attributable to intrinsic noise: stochastic changes in the molecular states of the promoter that have different abilities to transcribe. This is an important result because, although intrinsic noise is often considered as the main source of noise when modeling transcription imaging experiments, only a few studies have directly tested this hypothesis by assessing extrinsic versus intrinsic contributions using two-copy RNA imaging systems (67–69). These data suggest that for non-inducible genes, or inducible genes in the basal states, transcriptional fluctuations are mostly, if not entirely, attributable to the intrinsic dynamic of promoters.

In the case of HIV-1, direct RNA imaging has been used to characterize its stochastic promoter dynamics. The promoter has been shown to fluctuate across multiple time scales, from minutes to hours, a behavior interpreted as intrinsic stochastic switching between a single ON state and several OFF states (1,38,39). An interesting question is how TNFα and TSA modify these intrinsic promoter dynamics to increase transcription rates. TSA inhibits histone deacetylases (HDACs) and the most likely mechanism in this context is chromatin opening, where gene activation corresponds to an increased nucleosome mobility facilitated by TSA. This could be modeled as a promoter with multiple inactive states among which the longest corresponds to closed chromatin. TSA would then destabilize these long inactive periods, resulting in more frequent bursts consistent with our observations (Figure 5). In the case of TNFα, a previous study using a very similarly tagged HIV-1 system has shown the dependence of burst amplitude on the dose of TNFα (27). NF-kB has been shown to interact with multiple co-activators such as CBP and P/CAF (67). On HIV-1 specifically, activated NF-kB displaces a repressive HDAC1 complex and recruits P-TEFb to stimulate elongation (48,49,68). By these multiple effects, NF-kB could achieve the initial observed transcriptional pulse and subsequent increase in burst frequency, duration and amplitude (27 and Figure 5).

### Cellular heterogeneity accounts for variable responses to LRAs

In contrast to the basal state, extrinsic noise became readily detectable when the promoter was activated with TNFα or LRAs such as TSA, JQ1 or PMA-Ionomycin. It accounted for up to 40% of total noise and displayed differences depending on the inducer or the cell clone. While RNA imaging has been extensively used to study intrinsic noise in yeast, *Drosophila*, and human cells, studies investigating extrinsic noise sources remain scarce. Only two previous studies have used live-cell imaging of RNA in combination with a two-copy system to infer intrinsic and extrinsic noise contributions (67–69). Both studies used estrogen responsive genes, TFF1 and GREB1, and homozygous endogenous tagging with MS2 or PP7 sites to generate a homozygously tagged cell line that was imaged in presence of a sub-saturating amount of estrogen. Extrinsic noise was detected and accounted for the majority (68) or a minority of the total noise (69). Combined with our study, these results collectively suggest that, while intrinsic noise dominates expression variability in basal states, extrinsic noise becomes more apparent during transcriptional stimulation and is then responsible for a significant portion of expression heterogeneity.

While considering sources of extrinsic noise, it is conceptually useful to distinguish between distinct categories. Here, we separate cellular extrinsic noise, which encompasses factors that are specific to individual cells such as variation in transcription factor levels, metabolic state, or cell cycle stage, from environmental extrinsic noise, which refers to shared fluctuation that uniformly affects all cells in a population, such as changes in temperature, nutrient availability or extracellular signalling cues. This distinction has important implications for interpreting transcriptional heterogeneity and can be assessed by comparing gene copies in the same cell versus different cells. In the case of HIV-1, our results indicate that heterogeneity in the ability of single cells to respond to the activators play a major role in the variability of the viral transcriptional responses. Differences in the properties of individual cells were observed with all the compounds we tested (TNFα, TSA, JQ1, PMA-Ionomycin), which include both natural activators and synthetic drugs, suggesting that this may be a broadly general phenomenon. This raises a fundamental issue in the case of HIV-1 because cells failing to respond to activators remain in the body and provide the reservoir responsible for future viral rebounds. While ample evidence has shown that the intrinsic dynamics of the viral promoter can impact latency entry and exit (19,21,43), our results indicate that the heterogeneity of individual cells to sense and respond to viral activators also contribute to reservoir dynamics in patients, and that it largely contributes to the viral heterogeneous response to LRAs, which limits the efficacy of curing strategies such as shock-and-kill. Nailing down the causes of the heterogenous properties of single cells is thus an important issue for therapeutic approaches targeting the viral reservoir.

### Possible origins of cellular heterogeneity

The different sensitivity of single cells to viral activators may be fundamental for HIV-1 latency control and this raises the question of its origin. In the case of TNFα, it has been shown that signaling induces oscillations in NF-kB nuclear localization and that the heterogeneity of the response to TNFα is dependent on its dosage and stimulus duration (70). Interestingly, it has also been proposed that the response heterogeneity may arise from a variable abundance of TNFα receptors at the cell surface (71), a feature apparent in our cell system, and from negative and feed forward loops in the pathway that are designed to enhance heterogeneity (72). It has been suggested that the TNFα pathway has built-in characteristics that generate heterogeneities even in isogenic populations, which result from an evolutionary trade-off between the conflicting needs to respond efficiently to infection and avoid too much cell death (71). A heterogeneous response allows the pathway to rapidly kill infected cells while protecting many non-infected ones. Our results additionally indicate that noise due to the intrinsic dynamic of the target promoters also contributes significantly to the heterogeneity of the response to TNFα.

TSA inhibits histone deacetylases (HDACs), a family of four classes of enzymes comprising 18 members that catalyze the removal of acetyl groups from lysines, in particular on histone tails. TSA leads to chromatin decondensation and transcriptional activation, including for HIV-1 (73). The variability of the viral response to TSA showed a strong contribution of cellular extrinsic noise, indicating different abilities of individual cells to respond to the drug. This was observed in both the K11 and O1.4 clones, although with a stronger effect and earlier kinetics for K11. These differences in kinetics are likely due to clonal variability, which indirectly also underscores the heterogeneity in the properties of individual cells in the parental population. Given the role of noise for HIV-1 biology (21,32,43), it would be important to determine the exact cellular properties that make cells sense and respond differently to LRAs molecules like TSA. Interestingly, measurement of acetylated lysines in cells, the direct molecular consequence of HDAC inhibition, shows that TSA greatly increases the cell-to-cell variability in acetylated lysines, a visible and possibly long-lasting manifestation of the variable properties of individual cells detected by the two-copy imaging system. It is unclear how TSA could create this heterogeneity. The histone deacetylase HDAC1, a direct target of TSA, has been recently shown to be rapidly and strongly upregulated in response to TSA (74), suggesting a negative feedback loop in which cells respond to globally limit the effect of the inhibitor. Possibly, intrinsic noise in HDAC1 transcriptional response to TSA may translate into variable HDAC1 levels between cells, which becomes a global property of the cell that produces extrinsic noise for HDAC1 target genes. Similarly, it has been shown that TSA upregulates multidrug resistance genes (75,76), which may similarly convert an intrinsic noise into an extrinsic one, i.e. variable intracellular TSA concentrations. This would also yield heterogeneous cellular properties that can shape their viral response to the drug.

In the future, it will be interesting to test these hypotheses, for TSA and other drugs used as LRAs, with the aim of improving their therapeutic efficacy.

The existence of transcriptionally inactive latent subpopulations poses a significant barrier to the efficacy of shock-and-kill HIV-1 cure strategies. Enhancing the uniformity of cellular responses and increasing the proportion of activated cells could, therefore, improve therapeutic outcomes. While the intrinsic component of heterogeneity, driven by stochastic promoter switches, remains challenging to modulate, our results demonstrate that a substantial fraction of heterogeneity (up to 40%) stems from cell heterogeneity at the levels of signal transduction and phenotypic penetrance of drugs. These extrinsic components may be governed by molecular factors whose abundance and activity may be, at least in principle, amenable to modulation. Identifying these regulatory determinants is thus of critical importance to improve therapeutic strategies targeting the viral reservoir.

## Materials and Methods

### Cell culture and construction of CRISPR cassettes

HeLa cells were grown in Dulbecco’s modified Eagle’s Medium DMEM (Gibco #31966021) supplemented with 10% fetal bovine serum (Sigma Aldrich) and 1/100 U/ml Penicillin/streptomycin (Gibco #15140-122).

The HIV-1 reporter constructs were generated from the pIntro plasmid, which contains an optimized HIV-1 reporter with 128 MS2 loops (36). The pIntro plasmid also includes a 1500 bp stretch of human genomic DNA corresponding to a real HIV-1 integration site from a patient, located within the TBC1D5 gene and which corresponds to the right homology arm. Additionally, we cloned a 500 bp left homology arm upstream of the HIV-1 reporter. Downstream of this second homology arm, we incorporated a polylinker cloned two different selection markers for each repair cassette. In one cassette, we used a blasticidin selection marker under the control of the EF1α constitutive promoter. In the second cassette, we replaced the blasticidin marker with the hygromycin marker from the ptv-oct4 plasmid (Addgene #62351). Both selection markers were inserted in the reverse orientation relative to the HIV-1 LTR promoter. Plasmid sequences are available on request.

### Generation of HeLa CRISPR MCP-tdStayGold Clones

To insert the HIV-1 reporters into HeLa cells, we used one Cas9 guide RNA directing cleavage in between the two homology arms of our plasmids. The PAM and the guide binding sites were absent from the repair cassettes. The sequence of the sgRNA was 5’ TGGGTTACTCCTGAGTTAAC and it was cloned in pUC57-attbU6 sgRNA vector (77). Cas9-nickase expressing plasmid was provided from (Addgene #42335). Plasmids were transfected in a 3:3:1 ratio (repair cassette, U6 sgRNA, Cas9 Nickase) using Jet Prime (Polyplus #114-75), and cells were split after 24h and put under selection. Two weeks after transfection, individual clones were picked using cloning discs (Merck #F37847-0001) and cultured in 24 well plates under Hygromycin selection at 150 µg/ml final concentration following the first round of CRISPR transfection, and additionally under Blasticidin selection at 8 µg/ml following the second round of transfection. Clones were verified by genotyping and smiFISH after each round of transfection. Following the second CRISPR transfection, single cell clones were sorted to ensure clonal populations, the selected clones were again genotyped and verified by smiFISH. Selected HeLa clones were transduced with a lentivirus expressing NLS-MCP-tdStayGold (50), and FACS sorted to ensure stable expression.

### SmiFISH, immunofluorescence and fixed cell image acquisition

SmiFISH was performed as described previously (78). Briefly, 2.5x10^5^ cells were seeded on 22x22 mm glass coverslips. At 70% confluency, cells were fixed with 4% PFA in PBS for 20 min and permeabilized overnight with 70% ethanol at 4°C. Permeabilized cells were washed twice with PBS and incubated for 30 min at RT in a solution containing 15% formamide in 1xSSC. Unlabelled primary oligonucleotides were pre-hybridized with Cy3 or Cy5 labeled secondary oligonucleotides in a solution containing 100 mM NaCl, 50 mM tris- HCl, 10 mM MgCl_2_ (pH 7.9), for 3 min at 65°C and 10 minutes at RT to form a labeled duplex. Cells were hybridized overnight at 37°C with the duplex in the following mix: 15% Formamide, 0.34 mg/μl tRNA, 0.2 mg/ml RNase-free BSA, 2 mM VRC, 1xSSC and 10% dextran sulphate. Cells were washed 3 times for 45 minutes in 15% formamide 1xSSC, washed 3 times 1 minute with PBS, and the slides were mounted in a Vectashield containing DAPI (Vector Laboratories).

For immunofluorescence, cells were incubated in 0.2% Triton in PBS for 20 minutes. Cells were first blocked with 3% bovine serum albumin (BSA) in PBS for 30 minutes at room temperature. Following blocking, cells were incubated for 1h30 at room temperature with primary antibodies diluted in PBS containing 0.1% BSA; anti-NF-κB p65 (Cell Signaling Technology, #D14E12) at a 1:800 dilution, anti-TNFR1 (Proteintech, #60192-1-Ig) at a 1:100 dilution, and anti-acetylated lysine (Thermo Fisher Scientific, #MA1-2021) at a 1:500 dilution.

Cells were then washed 3x with 1xPBS, 10 minutes each. Following washes, cells were incubated with fluorescently labeled secondary antibodies: CY5-conjugated goat anti-rabbit IgG (Jackson ImmunoResearch, #111-175-144; 1:200) for NF-κB p65, CY5-conjugated donkey anti-mouse IgG (Jackson ImmunoResearch, #715-166-150; 1:200) for acetylated lysine, and CY3-conjugated donkey anti-mouse IgG (Jackson ImmunoResearch, #711-166-152; 1:200) for TNFR1.

Images were acquired using an upright fluorescent wide-field microscope (Zeiss Axioimager Z2 apotome). Images were taken using 63X objective (Apochromat 1.46 NA oil Corr), using Zen 3.10 software. Multi-dimensional acquisition was used to acquire 3D Z-stacks in 4 channels: DAPI, GFP, Cy3 and Cy5. The distance between each image in the stack was set to 0.3 micrometers. Maximum projection was performed to trans-form Z-stacks into 2D images. Figures were prepared with ImageJ, OMERO, Adobe Photoshop and Illustrator.

### Flow cytometry measurements and analysis

Hela K11 cells were plated in 10 cm dishes the day before the experiment. The next day cells were treated with TNFα (10 ng/ml) or TSA (400 nM) for the indicated time, fixed in 4% PFA and permeabilized with 0.2% TWEEN in 1xPBS (except for the TNFR1 labeling). The cells were incubated overnight at 4°C either with NF-kB p65 polyclonal antibody (1:100, Thermo Fisher Scientific, #51-0500), TNF Receptor 1 monoclonal antibody (TNFR1; 0.2 μg per 10^6^ cells,Proteintech, #60192-1-Ig) or Acetylated Lysine monoclonal antibody (1:200, Thermo Fisher Scientific, #MA1-2021). After washing, the cells were stained with the appropriate CY5 secondary antibody (1:200; Goat anti rabbit IgG, Jackson ImmunoResearch, #111-175-144 or Goat anti mouse IgG, Jackson ImmunoResearch, #115-175-166) and DAPI (5 μg/mL) for 1h at room temperature. The cells were analyzed with the NovoCyte Quanteon Flow Cytometer Systems 4 Lasers (Agilent) setting the filters R667 (ex/em, 637/667±30) for CY5 and V445 (ex/em, 405/445±45) for DAPI. Quantifications were done using FloJo v10.10.0. The intensity values were first assessed for normality and unimodality. A Mann–Whitney U test was performed between replicates to evaluate their similarity; upon confirmation of no significant differences, the replicates were pooled. From this aggregated dataset of 15826 cells, a subpopulation of approximately 7,000 values was randomly sampled to calculate the standard deviation and a dispersion metric (STD/mean). This resampling procedure was repeated 10,000 times to generate a distribution of dispersion values. A two-sample t-test was subsequently conducted to compare experimental groups and assess significant deviations from the null hypothesis.

### Live-cell image acquisition

Cells were plated on 35 mm glass-bottom imaging dishes “Fluorodish” (WPI, #FD35-100) and grown at 37°C to reach 70% confluence (usually within 48-72 hours). Prior to image acquisition, the cells were washed twice with PBS, and the culture medium was replaced with live-cell imaging medium (Fluorobrite DMEM; Thermo Fisher Scientific, #A1896701). The medium was supplemented with 1x GlutaMAX, 10% FBS, 500 µM ascorbic acid (Sigma, #A92902), 3 µM Rutin (Evrogen, #MCK02), and live-cell antifade reagent (Thermo Fisher Scientific, #P36974). Cells were then equilibrated in the microscopy incubation chamber for at least 20 minutes at 37°C and 5% CO2 before imaging.

Movie acquisition was conducted using an inverted Nikon microscope coupled to a Dragonfly spinning disk (Andor), equipped with a 100X Plan Apo 1.45 NA oil immersion objective. Excitation was achieved using a 488 nm laser, with an exposure time of 75 ms at 5% laser power in the GFP channel. Images were captured on an EMCCD iXon888 camera (1024 x 1024 pixels, effective pixel size: 0.121 µm) in multi-position acquisition mode. Z-stacks consisting of 13-15 optical slices, each separated by 0.6 µm, were acquired every 1 minute for a total of 13.5 hours. The perfect focus system (PFS) was employed, and 10 to 12 different stage positions were imaged during each session. No signs of cellular stress observed during the imaging period and cells divided normally while imaged.

For drug treatments, TNFα, TSA, JQ1, PMA-Ionomycin were introduced to the plate on-stage through a small opening in the plate cover, after 1 hour of imaging in basal conditions. The drug of interest, dissolved in an equal volume of Fluorobrite, was added dropwise to the medium.

### RNA quantification from smiFISH image data

SmiFISH analysis was performed using a friendly user application called “SmallFISH”, which is accessible through GitHub: https://github.com/2Echoes/small_fish_gui. It operates by integrating cell segmentation using Cellpose, a deep learning-based algorithm (79), with single molecule detection through BigFISH, which employs a noise-robust local maxima filter for precise detection (80). The software then extracts a variety of features for each cell (number of RNA, transcription site intensities), enabling statistical analysis across large batches of cells.

### Live cell quantifications

Quantification of transcription sites intensities from raw microscopy time-lapse imaging data was performed in several steps.

#### Pre-Processing

We performed maximum intensity projections along both the Z-axis and the temporal axis for each field of view in order to outline the overall spatial footprint of each nucleus. Nuclear segmentation was subsequently carried out at each time point using an automated pipeline based on Cellpose (79). This pipeline also enabled cropping of individual nuclei and temporal alignment of their centroids to generate motion-corrected time-lapse image series for all nuclei within a field of view. Photobleaching was quantified by measuring the sum of pixel intensities within nuclear masks and comparing these values to intensities measured in background regions. For three-dimensional image stacks exhibiting greater than 20% photobleaching, correction was performed using the bleach correction plugin in Napari. Missing frames were imputed by calculating the mean image from adjacent time points (t−1 and t+1).

#### Spot detection and transcription site quantification

Spot detection thresholds and clustering parameters were empirically optimized for each cell from a sample of frames spaced across the temporal dimension. Using the Spot and Cluster Detection module of liveQuant (https://github.com/rachel-kt/liveQuant), spots were detected across all frames. The detected spots were then used to construct a reference spot representing the fluorescence signal of a single RNA molecule. This reference model was then used to decompose detected clusters into estimated counts of individual RNA molecules, as done in bigFISH (80). Transcription site (TS) positions were initially detected in each frame by applying a computational blurring technique designed to attenuate single-molecule signals while preserving the transcription site signal. These generated tracks that were then manually corrected and used to obtain high confidence transcription site signals.

### Data preparation and noise analysis

#### Total variance decomposition to obtain intracellular and environmental extrinsic noise components

Consider a two reporter system with two transcription sites like in Figure 7F. The two sites function independently but both collect signals from the environment. We suppose that the sites are otherwise equivalent and contain replicas of the same promoter and cis-regulatory regions. In order to quantify extrinsic noise, we consider the covariance of the RNAs produced by two different sites. We consider two situations: i) paired sites, when the two sites are chosen in the same cell and ii) unpaired sites, when the two sites are chosen in different cells. By doing so we probe two types of extrinsic noise: intracellular and environmental. Intracellular sources affect each cell differently, while environmental sources affect groups of cells identically. These latter sources include changes in the external environment, such as the presence or absence of different cell types and their densities, mechanical stress, and signaling molecules or drugs. Intracellular sources involve heterogeneity in transcription factors levels, metabolism, signaling molecules, as well as cell-cycle stages.

Adapting reasonings from (7, 81, 82) we formalize the problem as follows:

1. The variable *Z^cell^* represents the cellular internal environment and the variable *Z^env^* represents the external environment shared by several cells. Let *x* be the measured gene product (nascent, or total RNA).
2. The extrinsic noise is represented by the conditional expectation *x̃_extr_* = 𝔼[*x*|*Z^cell^*, *Z^env^*].
3. The environmental extrinsic noise is *x̃_env_* = 𝔼[*x*|*Z^env^*].
4. The cellular extrinsic noise is *x̃_cell_* = *x̃_extr_* − *x̃_env_*.
5. Finally, the intrinsic noise is *x̃_intr_* = *x* − *x̃_extr_*

The tower property of the conditional expectation, also known as the law of total variance leads to

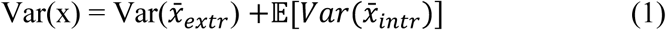

Furthermore,

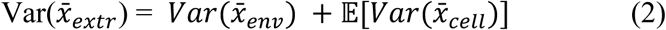

Adapting results from (7) (82) to our experimental design, the covariance of unpaired sites can be used to estimate *Var*(*x̃_env_*), whereas the covariance of paired sites provides Var(*x̃_extr_*.

According to (1) and (2), the variance of single site signal decomposes in three components: intrinsic, intracellular extrinsic and environmental extrinsic noise.

#### Fixed Cell Experiments

The mean counts of nascent RNA per transcription site and released RNA per nucleus for fixed cell data were calculated and presented as histograms, which were plotted using the Seaborn Python data visualization package. Additionally, correlation plots between the two transcription sites for each condition were generated for each clone, providing further insight into the relationship between the transcription dynamics. The extrinsic and intrinsic noise were computed using the law of total variance (see Equations (1) and (2)). Using (7, 82) the contributions of extrinsic and intrinsic noise to total noise were estimated using the following formulas:

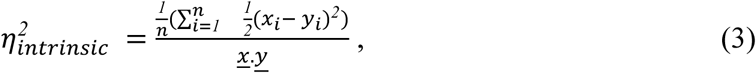

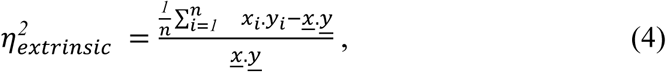

where *x_i_*, *y_i_* are values of mRNA numbers for pairs of transcription sites, and *x̃*, *ỹ* are the respective means. In (3) the sum is over both true pairs (same cell TS) and simulated random pairing (TS from different cells). When in (4) the sum is over true pairs, *η^2^_extrinsic_* estimates the contribution of cellular and environmental noise, whereas a sum over unpaired sites in (4) estimates only the environmental noise, where n is the number of pairs.

#### Live Cell Experiments

Each dataset consists of time-resolved traces of transcription site (TS) RNA copy number, from multiple cells under distinct experimental conditions, such as basal or drug-treated. A custom Python script was employed to compute cumulative noise metrics using a time cumulative approach that incorporates data from preceding time points. This method enables the quantification of intrinsic, extrinsic, and total noise components as they evolve over time. Formulas (3) and (4) were thus modified to account for the temporal evolution of the noise as follows

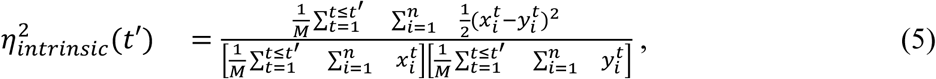

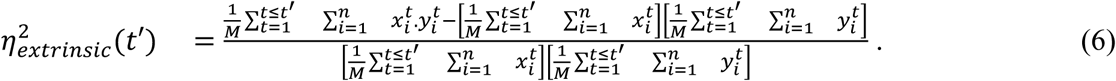

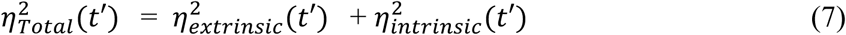

Here, 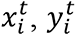 represent the mRNA values from the two transcription sites at a particular timepoint t.

This cumulative approach enables a progressively refined assessment of noise contributions, capturing temporal trends in transcriptional variability and avoiding the limitations of analyzing isolated time points, as instantaneous estimators are less robust and exhibit greater variability. To demonstrate this, instantaneous noise metrics were also calculated by considering data from individual time points. A moving average (window size=5) of these instantaneous noise values was applied to reduce the impact of outliers, as depicted in Figure 7 plots C and D, along with an overlay of the cumulative values of true pairs. The code for computing the instantaneous and cumulative noise is available on GitHub: https://github.com/rachel-kt/Extrinsic_Noise

For the autocorrelation analysis, the auto-correlation function of each trace and the cross-correlation functions between two true paired (same cell TS) or unpaired traces (TS from different cells) were defined as

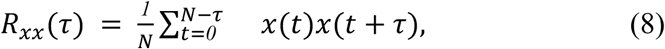

and were calculated using the inverse Fast Fourier Transform as

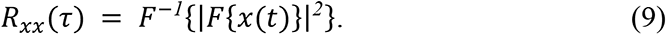

The average correlation functions were then plotted for each condition and clone. These analyses were performed using custom Python scripts, employing the SciPy and NumPy libraries. To eliminate bias in selecting the “first site”, the RNA trajectories were shuffled. Additionally, for computing the correlation function of unpaired traces, transcription sites were shuffled such that two randomly selected transcription sites were never from the same nucleus. A bootstrap of 50 random permutations of pairs of transcription site trajectories equal to the total number of cells were performed to obtain an average cross-correlation function for unpaired traces (Figure 6A and 6B).

The autocorrelation functions contain information about the timescales of the noise, which can be extracted through multi-exponential regression. Specifically, we employ the following parametric model:

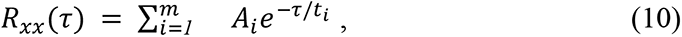

where *t_i_* > *0* are timescales and *A_i_* are other parameters.

The number of exponentials is determined by a parsimony criterion. Starting with *m*=1, we progressively increase *m* until the least squares error of the regression becomes sufficiently small, while ensuring that the uncertainty in the timescale parameters remains limited, in order to avoid overfitting. The uncertainty error is estimated by computing a set of optimal and suboptimal solutions to the regression problem, and taking the minimum and maximum parameter values within this set.

In Equation (9), we assume that the exponents are real. However, cases involving complex exponents and oscillatory behavior of the autocorrelation function are also encompassed by the same formula, when *A_i_* and *t_i_* are complex parameters. Since the number of real parameters doubles in such cases (each complex parameter has real and imaginary parts), we did not consider them to avoid overfitting.

## Data Availability

The raw data for live and fixed imaging are provided in the Supplementary Spreadsheet and the code used is available on Github as indicated. All data needed to evaluate the conclusions in the paper are present in the manuscript and the Supplementary Materials. All the materials used in the study is available upon request.

## Acknowledgements

We thank Florian Müller and Thomas Walter for stimulating discussions and critical review of this work. We acknowledge and thank members of the MRI imaging facility at IGH and CRBM. MRI is part of the national infrastructure France-BioImaging supported by the French National Research Agency (ANR-10-INBS-04, ‘Investissement d’Avenir’ program). R.T was supported by the Labex EpiGenMed (LabMuse) and Sidaction. H.K was supported by (CNRS-Liban/UM) and Sidaction. F.M was supported by ANRS-MIE. This work was supported by grant ECTZ176246 from ANRS-MIE. R.T developed the tools for noise analysis, for live-cell image analysis, and analyzed the data. H.K constructed and characterized the cell lines and performed the fixed and live-cell imaging experiments. F.M., O.P. and M.P. helped to analyze the live-cell imaging data. F.M. performed the FACS analyses.

K.Z. co-supervised H.K. O.R. and E.B. conceived the study and supervised the project. R.T., H.K., O.R. and E.B. generated the figures and wrote the paper.

## List of Supplementary Materials

### Supplementary Spreadsheets

**Supplementary Spreadsheet 1:** RNA count per cell computed from the fixed cell experiments.

**Supplementary Spreadsheet 2:** RNA count per transcription site computed from the live cell experiments.

## Supplementary Videos

**Video S1**: Time-lapse movie showing a cell of the K11 clone under basal conditions, acquired over 10 hours with images captured every minute. A maximum intensity projection is shown.

**Video S2**: Time-lapse movie showing a cell of the K11 clone treated with 10ng/ml TNF-α, acquired over 10 hours with images captured every minute. A maximum intensity projection is shown.

**Video S3**: Time-lapse movie showing a cell of the K11 clone treated with 400 nM TSA, acquired over 10 hours with images captured every minute. A maximum intensity projection is shown.

**Video S4**: Time-lapse movie showing a cell of the K11 clone treated with PMA-Ionomycine, acquired over 10 hours with images captured every minute. A maximum intensity projection is shown.

**Video S5**: Time-lapse movie showing a cell of the K11 clone treated with 1 µm JQ1, acquired over 10 hours with images captured every minute. A maximum intensity projection is shown.

**Video S6**: Time-lapse movie showing a cell of the O1.4 clone under basal conditions, acquired over 10 hours with images captured every minute. A maximum intensity projection is shown.

**Video S7**: Time-lapse movie showing a cell of the O1.4 clone treated with 400 nM TSA, acquired over 10 hours with images captured every minute. A maximum intensity projection is shown.

## Supplementary Figures

**Supplementary Figure 1.**
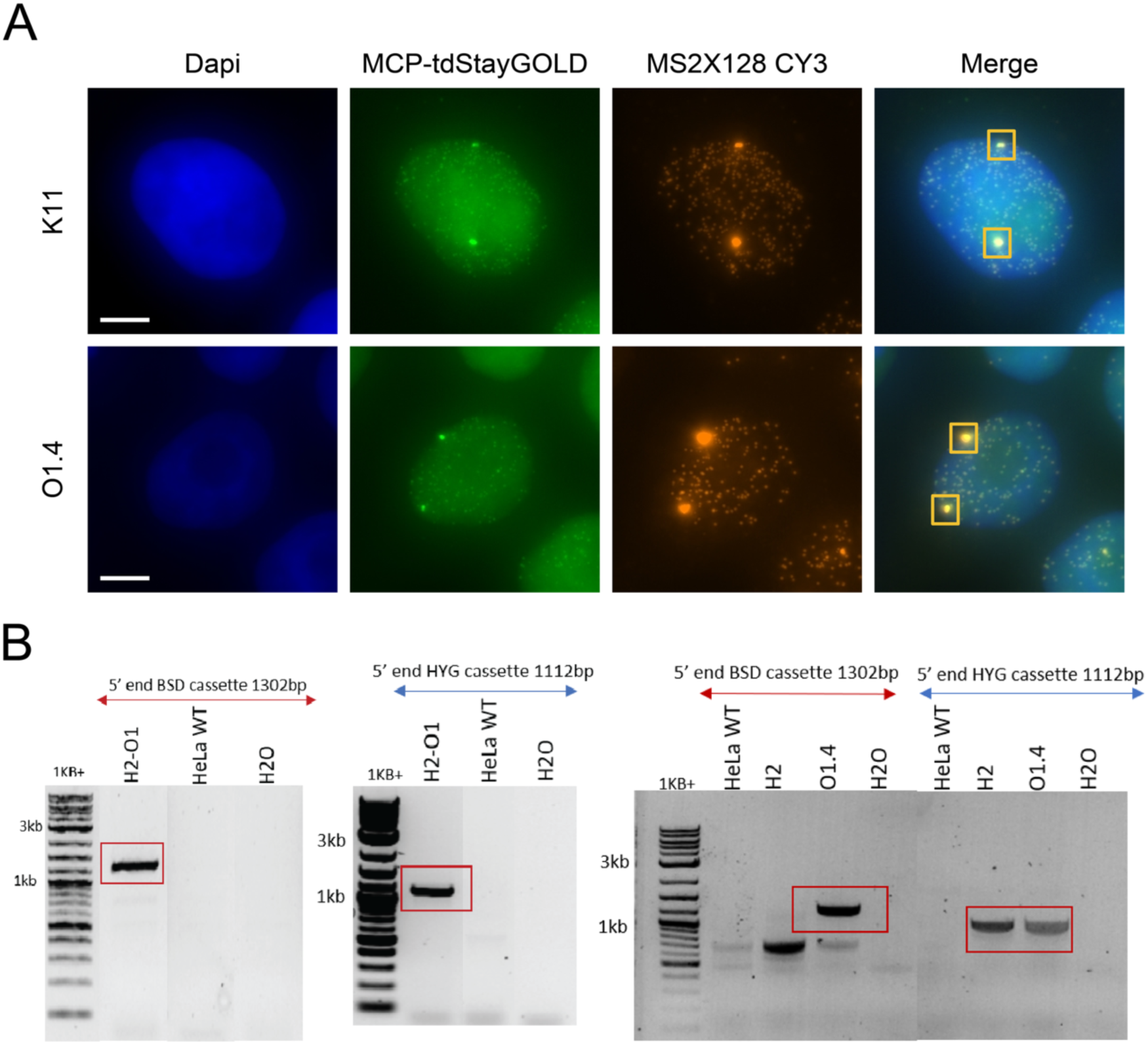
Characterization of the two copy HIV-1 CRISPR clones. **A.** Micrographs of K11 (upper panel) and O1.4 (lower panel) cells, 8 hours after treatment with TSA. From left to right: DAPI (blue) stains nuclear DNA; MCP-tdStayGold (green) marks nuclear MCP signal; and Cy3 (red) visualizes smiFISH targeting the MS2 stem-loops. Transcription sites are highlighted with yellow boxes. Scale bar: 5 µm. **B.** The image displays three agarose gels loaded with PCR products targeting the 5’ end of the integration sites for two repair cassettes, following amplification of genomic DNA extracts. The expected sizes for the DNA products corresponding to the cassettes containing the blasticidin and hygromycin genes are 1302 bp and 1113 bp, respectively. The H2 clone exhibits only the band for the first CRISPR, while clones O1 and O1.4 display the expected bands for both repair cassettes. A HeLa wild-type DNA extract was used as a negative control, with no specific bands detected, and H2O served as a PCR negative control. BSD refers to blasticidine and HYG refers to hygromycine.

**Supplementary Figure 2.**
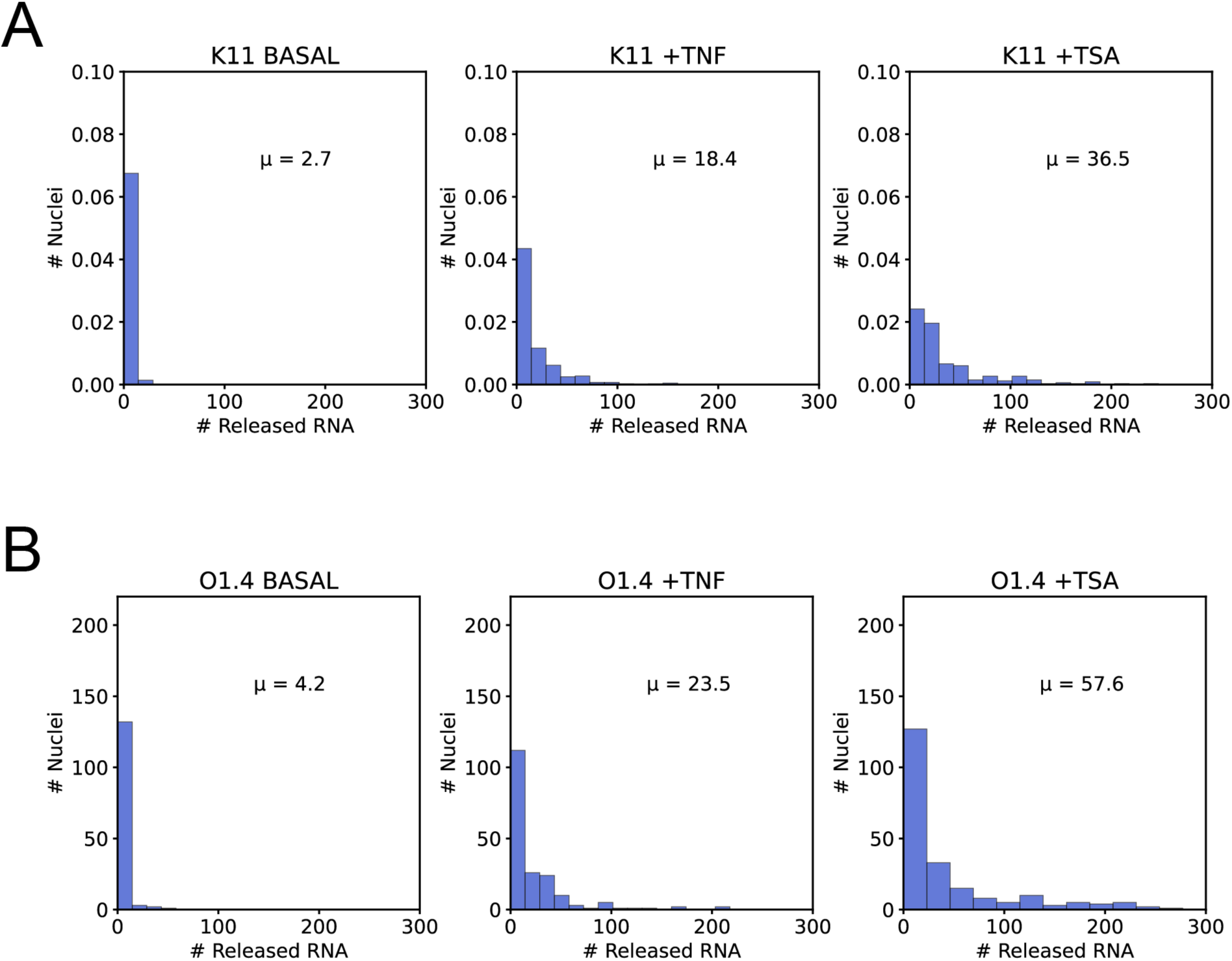
Distribution of the number of released RNA molecules per nucleus across conditions in K11 and O1.4 cell lines. **A-B**. Distribution of released RNA counts per nucleus for K11 (top row) and O1.4 cells (bottom row), under three conditions: basal, TNFα, and TSA (from left to right). The x-axis represents the number of released RNA molecules per nucleus, while the y-axis indicates the number of nuclei with that respective RNA count. The mean RNA count (μ) for each condition is shown in each plot.

**Supplementary Figure 3.**
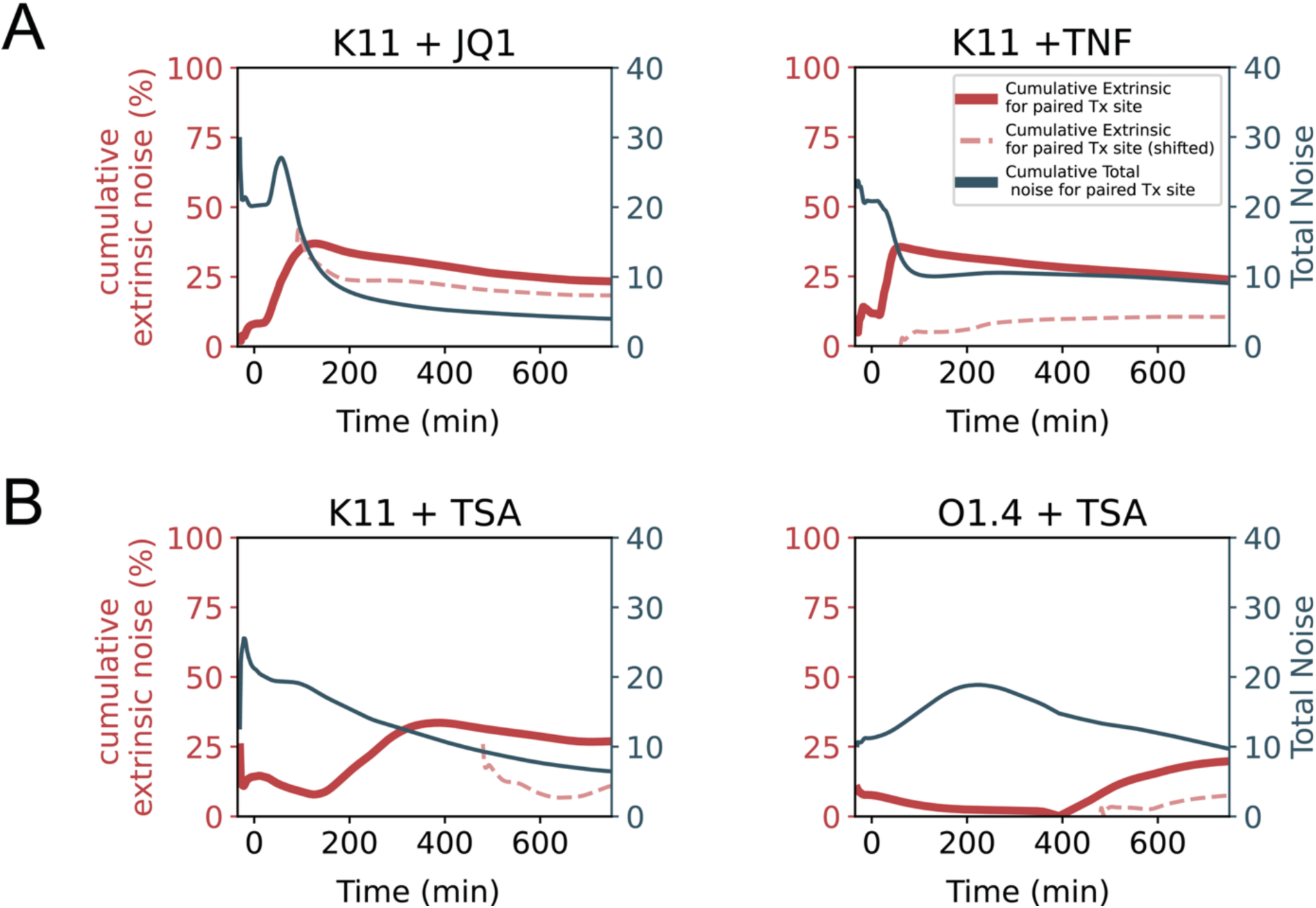
Cumulative extrinsic noise analysis over time for clones K11 and O1.4. **A-B.** The plots display the cumulative extrinsic noise for clone K11 under TSA and TNF treatment (A), and K11 under JQ1 and O1.4 under TSA (B). The bold red line represents the cumulative extrinsic noise, measured as a percentage of the total noise, while the solid blue line denotes the total noise. The dotted red line shows the cumulative extrinsic noise calculated starting at 60 minutes for TNFα treatment, 480 minutes for TSA treatment and 90 min for JQ1. The left y-axis corresponds to cumulative extrinsic noise in percentage, and the right y-axis represents total noise.

**Supplementary Figure 4.**
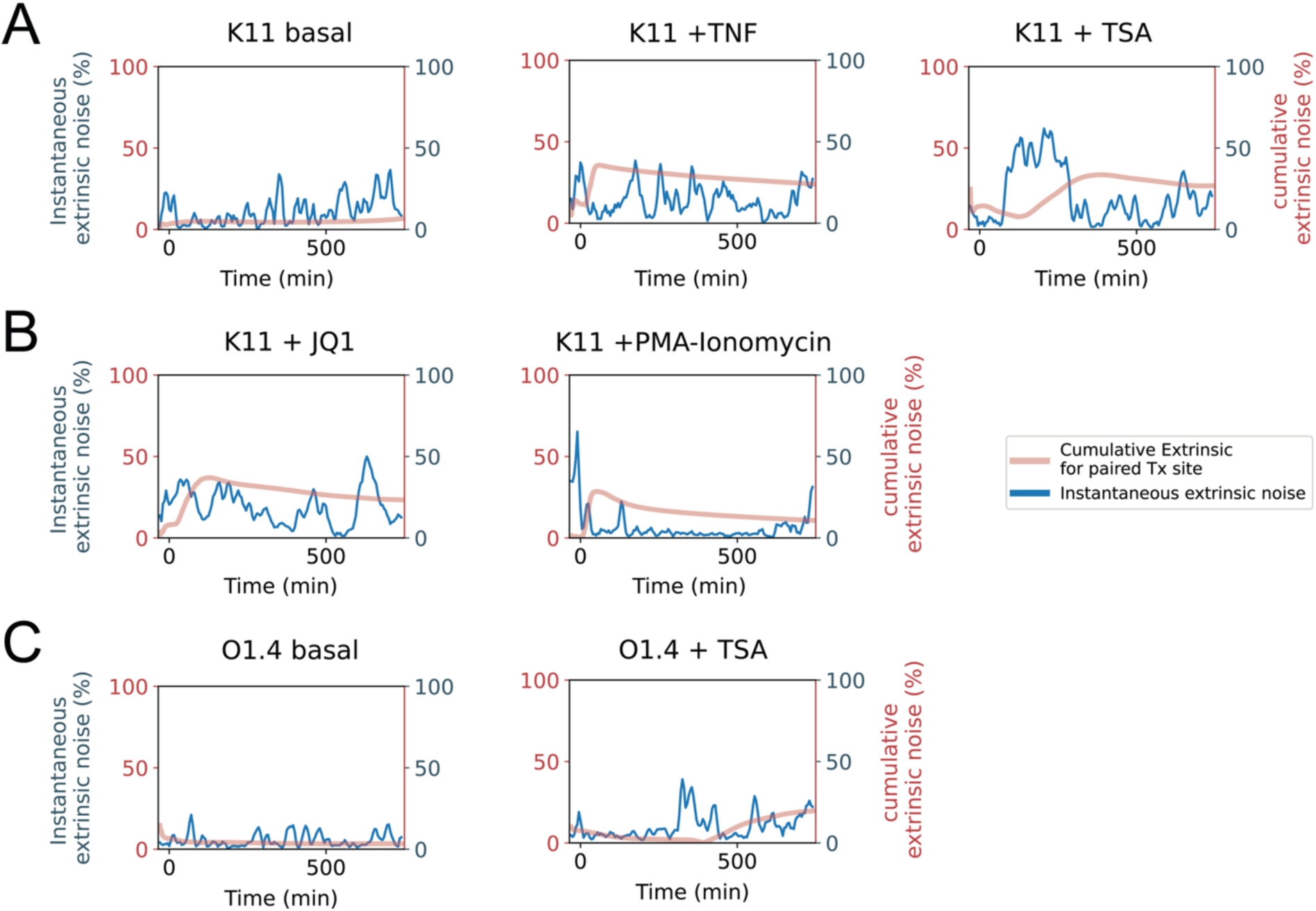
Instantaneous vs. cumulative extrinsic noise in clones K11 and O1.4. **A-C.** Plots display the cumulative extrinsic noise and instantaneous extrinsic noise for paires transcription sites, for clone K11 (A and B) and O1.4 (C), measured as a percentage of total noise and calculated at each time point under basal condition and after the indicated treatments. The bold red lines represent the cumulative extrinsic noise for paired transcription sites (as a percentage of total noise), while the blue line denotes the instantaneous extrinsic noise for unpaired transcription sites (also measured as a percentage of total noise).

**Supplementary Figure 5.**
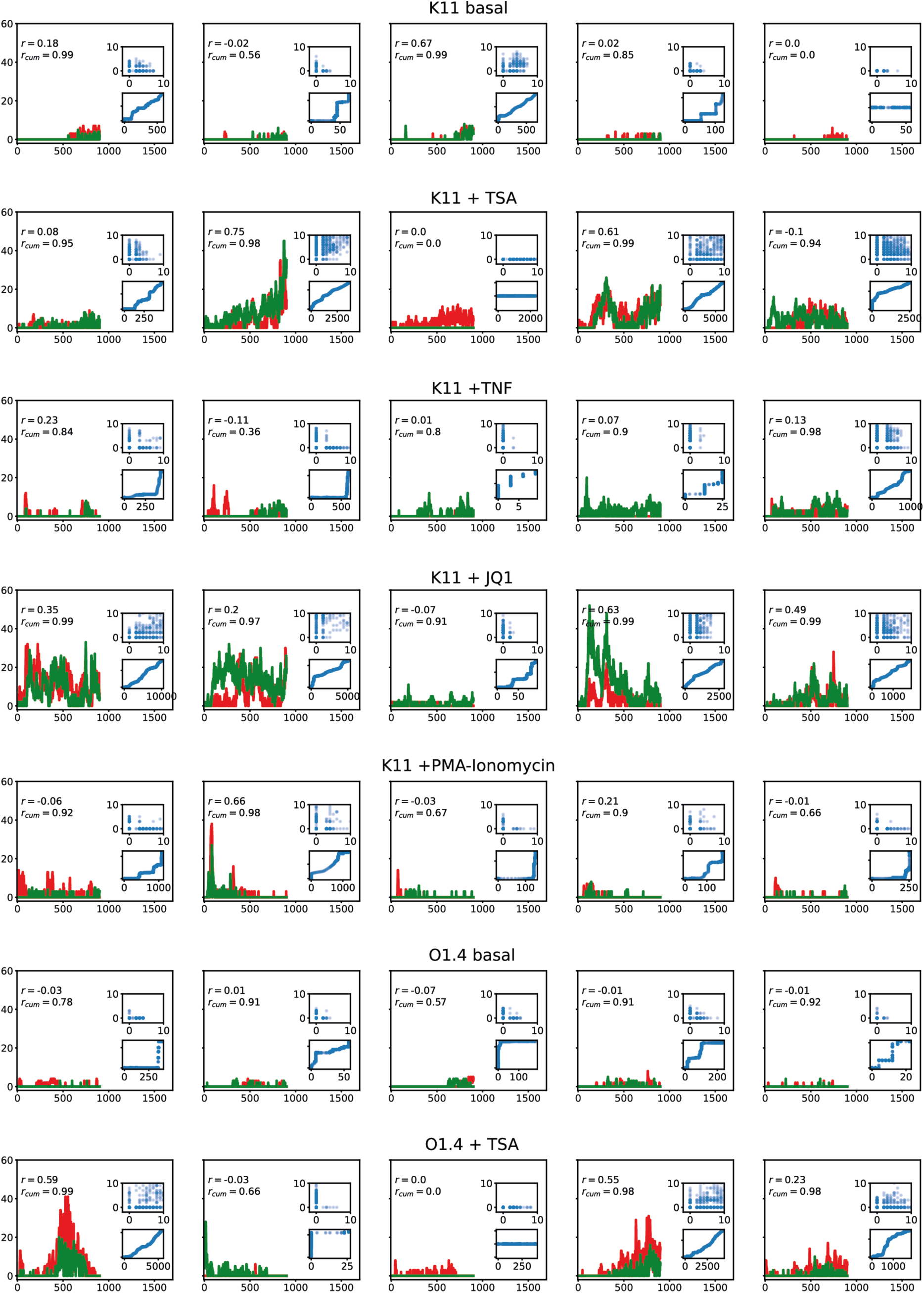
Examples of nascent RNA transcription at dual transcription sites in single cells. The plots show the nascent RNA trajectories in time for the two transcription sites (red and green) in single cells. The top most inset is a scatter plot of the nascent RNA numbers from the two transcription sites at each time point. Zero nascent RNA means the transcription site is ’OFF’. The bottom inset displays the integrated sum of the nascent RNA numbers at the two transcription sites. The Pearson correlation coefficient of transcription site intensities (r), is displayed at the top left corner of each plot. The Pearson correlation coefficient of the cumulative transcription site intensities (r_cum_), indicating the linear relationship between the two transcription sites, is displayed at the top-left corner of each plot.

## References

1. Chubb JR, Trcek T, Shenoy SM, Singer RH. Transcriptional Pulsing of a Developmental Gene. Curr Biol. 2006 May;16(10):1018–25.

2. Beckman WF, Jiménez MÁL, Verschure PJ. Transcription bursting and epigenetic plasticity: an updated view. Epigenetics Commun.. 2021 Dec;1(1). Available from: 10.1186/s43682-021-00007-1

3. Raj A, Rifkin SA, Andersen E, van Oudenaarden A. Variability in gene expression underlies incomplete penetrance. Nature. 2010 Feb 18;463(7283):913–8.

4. Weinberger LS, Burnett JC, Toettcher JE, Arkin AP, Schaffer DV. Stochastic gene expression in a lentiviral positive-feedback loop: HIV-1 Tat fluctuations drive phenotypic diversity. Cell. 2005 July 29;122(2):169–82.

5. Wernet MF, Mazzoni EO, Çelik A, Duncan DM, Duncan I, Desplan C. Stochastic spineless expression creates the retinal mosaic for colour vision. Nature. 2006 Mar 9;440(7081):174–80.

6. Urban EA, Johnston RJ. Buffering and Amplifying Transcriptional Noise During Cell Fate Specification. Front Genet. 2018;9:591.

7. Elowitz MB, Levine AJ, Siggia ED, Swain PS. Stochastic Gene Expression in a Single Cell. Science. 2002 Aug;297(5584):1183–6.

8. Lei X, Tian W, Zhu H, Chen T, Ao P. Biological Sources of Intrinsic and Extrinsic Noise in cI Expression of Lysogenic Phage Lambda. Sci Rep. 2015 Sept;5(1). Available from: 10.1038/srep13597

9. Swain PS, Elowitz MB, Siggia ED. Intrinsic and extrinsic contributions to stochasticity in gene expression. Proc Natl Acad Sci. 2002 Oct;99(20):12795–800.

10. Tunnacliffe E, Chubb JR. What is a transcriptional burst? Trends Genet. 2020;36(4):288–97.

11. Nicolas D, Phillips NE, Naef F. What shapes eukaryotic transcriptional bursting? Mol Biosyst. 2017;13(7):1280–90.

12. Urban EA, Johnston RJ. Buffering and Amplifying Transcriptional Noise During Cell Fate Specification. Front Genet. 2018 Nov;9. Available from: 10.3389/fgene.2018.00591

13. Raser JM, O’Shea EK. Noise in Gene Expression: Origins, Consequences, and Control. Science. 2005 Sept;309(5743):2010–3.

14. Krebs AR, Imanci D, Hoerner L, Gaidatzis D, Burger L, Schübeler D. Genome-wide Single-Molecule Footprinting Reveals High RNA Polymerase II Turnover at Paused Promoters. Mol Cell. 2017 Aug;67(3):411–422.e4.

15. Brown CR, Mao C, Falkovskaia E, Jurica MS, Boeger H. Linking Stochastic Fluctuations in Chromatin Structure and Gene Expression. PLOS Biol. 2013 Aug 6;11(8):e1001621.

16. Volfson D, Marciniak J, Blake WJ, Ostroff N, Tsimring LS, Hasty J. Origins of extrinsic variability in eukaryotic gene expression. Nature. 2006 Feb;439(7078):861–4.

17. Johnston IG, Gaal B, Neves RP das, Enver T, Iborra FJ, Jones NS. Mitochondrial Variability as a Source of Extrinsic Cellular Noise. PLOS Comput Biol. 2012 Mar 8;8(3):e1002416.

18. Cole JA, Luthey-Schulten Z. Careful accounting of extrinsic noise in protein expression reveals correlations among its sources. Phys Rev E. 2017 June 27;95(6):062418.

19. Damour A, Slaninova V, Radulescu O, Bertrand E, Basyuk E. Transcriptional Stochasticity as a Key Aspect of HIV-1 Latency. Viruses. 2023 Sept;15(9):1969.

20. Sherman MS, Lorenz K, Lanier MH, Cohen BA. Cell-to-cell variability in the propensity to transcribe explains correlated fluctuations in gene expression. Cell Syst. 2015;1(5):315–25.

21. Weinberger AD, Weinberger LS. Stochastic Fate Selection in HIV-Infected Patients. Cell. 2013 Oct;155(3):497–9.

22. Siliciano JD, Siliciano RF. In Vivo Dynamics of the Latent Reservoir for HIV-1: New Insights and Implications for Cure. Annu Rev Pathol. 2022 Jan 24;17:271–94.

23. Siliciano JD, Siliciano RF. HIV cure: The daunting scale of the problem. Science. 2024 Feb 16;383(6684):703–5.

24. Harrich D, Garcia J, Wu F, Mitsuyasu R, Gonazalez J, Gaynor R. Role of SP1-binding domains in in vivo transcriptional regulation of the human immunodeficiency virus type 1 long terminal repeat. J Virol. 1989 June;63(6):2585–91.

25. Perkins ND, Edwards NL, Duckett CS, Agranoff AB, Schmid RM, Nabel GJ. A cooperative interaction between NF-κ B and Sp1 is required for HIV-1 enhancer activation. EMBO J. 1993 Sept;12(9):3551–8.

26. Wong VC, Bass VL, Bullock ME, Chavali AK, Lee REC, Mothes W, et al. NF-κB-Chromatin Interactions Drive Diverse Phenotypes by Modulating Transcriptional Noise. Cell Rep. 2018 Jan;22(3):585–99.

27. Zambrano S, Loffreda A, Carelli E, Stefanelli G, Colombo F, Bertrand E, et al. First Responders Shape a Prompt and Sharp NF-κB-Mediated Transcriptional Response to TNF-α. iScience. 2020 Sept;23(9):101529.

28. Cary DC, Fujinaga K, Peterlin BM. Molecular mechanisms of HIV latency. J Clin Invest. 2016 Jan;126(2):448–54.

29. Schulze-Gahmen U, Echeverria I, Stjepanovic G, Bai Y, Lu H, Schneidman-Duhovny D, et al. Insights into HIV-1 proviral transcription from integrative structure and dynamics of the Tat:AFF4:P-TEFb:TAR complex. eLife. 2016 Oct;5. Available from: 10.7554/eLife.15910

30. Gu J, Babayeva ND, Suwa Y, Baranovskiy AG, Price DH, Tahirov TH. Crystal structure of HIV-1 Tat complexed with human P-TEFb and AFF4. Cell Cycle. 2014 Apr;13(11):1788–97.

31. Hansen MMK, Martin B, Weinberger LS. HIV Latency: Stochastic across Multiple Scales. Cell Host Microbe. 2019 Dec;26(6):703–5.

32. Singh A, Razooky B, Cox CD, Simpson ML, Weinberger LS. Transcriptional Bursting from the HIV-1 Promoter Is a Significant Source of Stochastic Noise in HIV-1 Gene Expression. Biophys J. 2010 Apr;98(8):L32–4.

33. Singh A, Razooky BS, Dar RD, Weinberger LS. Dynamics of protein noise can distinguish between alternate sources of gene-expression variability. Mol Syst Biol. 2012 Jan;8(1). Available from: 10.1038/msb.2012.38

34. Weinberger LS. A minimal fate-selection switch. Curr Opin Cell Biol. 2015 Dec;37:111–8.

35. Weinberger LS, Shenk T. An HIV Feedback Resistor: Auto-Regulatory Circuit Deactivator and Noise Buffer. Aitchison JD, editor. PLoS Biol. 2006 Dec;5(1):e9.

36. Tantale K, Mueller F, Kozulic-Pirher A, Lesne A, Victor JM, Robert MC, et al. A single-molecule view of transcription reveals convoys of RNA polymerases and multi-scale bursting. Nat Commun. 2016 July;7(1). Available from: 10.1038/ncomms12248

37. Tantale K, Garcia-Oliver E, Robert MC, L’Hostis A, Yang Y, Tsanov N, et al. Stochastic pausing at latent HIV-1 promoters generates transcriptional bursting. Nat Commun. 2021 July 23;12(1):4503.

38. Pierson T, McArthur J, Siliciano RF. Reservoirs for HIV-1: Mechanisms for Viral Persistence in the Presence of Antiviral Immune Responses and Antiretroviral Therapy. Annu Rev Immunol. 2000 Apr;18(1):665–708.

39. Lorenzo-Redondo R, Fryer HR, Bedford T, Kim EY, Archer J, Kosakovsky Pond SL, et al. Persistent HIV-1 replication maintains the tissue reservoir during therapy. Nature. 2016 Jan;530(7588):51–6.

40. Kim Y, Anderson JL, Lewin SR. Getting the “Kill” into “Shock and Kill”: Strategies to Eliminate Latent HIV. Cell Host Microbe. 2018 Jan;23(1):14–26.

41. Lichterfeld M. Reactivation of latent HIV moves shock-and-kill treatments forward. Nature. 2020 Jan;578(7793):42–3.

42. Razooky BS, Pai A, Aull K, Rouzine IM, Weinberger LS. A hardwired HIV latency program. Cell. 2015;160(5):990–1001.

43. Weinberger LS, Burnett JC, Toettcher JE, Arkin AP, Schaffer DV. Stochastic Gene Expression in a Lentiviral Positive-Feedback Loop: HIV-1 Tat Fluctuations Drive Phenotypic Diversity. Cell. 2005 July;122(2):169–82.

44. Ho YC, Shan L, Hosmane NN, Wang J, Laskey SB, Rosenbloom DIS, et al. Replication-Competent Noninduced Proviruses in the Latent Reservoir Increase Barrier to HIV-1 Cure. Cell. 2013 Oct;155(3):540–51.

45. Jordan A. The site of HIV-1 integration in the human genome determines basal transcriptional activity and response to Tat transactivation. EMBO J. 2001 Apr;20(7):1726–38.

46. Bertrand E, Chartrand P, Schaefer M, Shenoy SM, Singer RH, Long RM. Localization of ASH1 mRNA particles in living yeast. Mol Cell. 1998 Oct;2(4):437–45.

47. Bellec M, Chen R, Dhayni J, Trullo A, Avinens D, Karaki H, et al. Boosting the toolbox for live imaging of translation. RNA. 2024 July;30(10):1374–94.

48. Williams SA, Chen LF, Kwon H, Ruiz-Jarabo CM, Verdin E, Greene WC. NF-κB p50 promotes HIV latency through HDAC recruitment and repression of transcriptional initiation. EMBO J. 2005 Dec;25(1):139–49.

49. Williams SA, Kwon H, Chen LF, Greene WC. Sustained induction of NF-kappa B is required for efficient expression of latent human immunodeficiency virus type 1. J Virol. 2007 June;81(11):6043–56.

50. Duh EJ, Maury WJ, Folks TM, Fauci AS, Rabson AB. Tumor necrosis factor alpha activates human immunodeficiency virus type 1 through induction of nuclear factor binding to the NF-kappa B sites in the long terminal repeat. Proc Natl Acad Sci U S A. 1989 Aug;86(15):5974–8.

51. Rao J, Bhattacharya D, Banerjee B, Sarin A, Shivashankar GV. Trichostatin-A induces differential changes in histone protein dynamics and expression in HeLa cells. Biochem Biophys Res Commun. 2007 Nov 16;363(2):263–8.

52. Bass VL, Wong VC, Bullock ME, Gaudet S, Miller-Jensen K. TNF stimulation primarily modulates transcriptional burst size of NF-κB-regulated genes. Mol Syst Biol. 2021 July;17(7). Available from: 10.15252/msb.202010127

53. Finnegan A, Roebuck KA, Nakai BE, Gu DS, Rabbi MF, Song S, et al. IL-10 cooperates with TNF-alpha to activate HIV-1 from latently and acutely infected cells of monocyte/macrophage lineage. J Immunol. 1996 Jan;156(2):841–51.

54. Kiernan RE. HIV-1 Tat transcriptional activity is regulated by acetylation. EMBO J. 1999 Nov;18(21):6106–18.

55. Li Z, Guo J, Wu Y, Zhou Q. The BET bromodomain inhibitor JQ1 activates HIV latency through antagonizing Brd4 inhibition of Tat-transactivation. Nucleic Acids Res. 2013 Jan 1;41(1):277–87.

56. Touraine JL, Hadden JW, Touraine F, Hadden EM, Estensen R, Good RA. Phorbol myristate acetate: a mitogen selective for a T-lymphocyte subpopulation. J Exp Med. 1977 Feb 1;145(2):460–5.

57. Toeplitz BK, Cohen AI, Funke PT, Parker WL, Gougoutas JZ. Structure of ionomycin - a novel diacidic polyether antibiotic having high affinity for calcium ions. J Am Chem Soc. 1979 June 1;101(12):3344–53.

58. Chen G, Goeddel DV. TNF-R1 signaling: a beautiful pathway. Science. 2002 May 31;296(5573):1634–5.

59. Aggarwal BB. Signalling pathways of the TNF superfamily: a double-edged sword. Nat Rev Immunol. 2003 Sept;3(9):745–56.

60. Liu ZG. Molecular mechanism of TNF signaling and beyond. Cell Res. 2005 Jan;15(1):24–7.

61. de Ruijter AJM, van Gennip AH, Caron HN, Kemp S, van Kuilenburg ABP. Histone deacetylases (HDACs): characterization of the classical HDAC family. Biochem J. 2003 Mar 15;370(Pt 3):737–49.

62. Ailenberg M, Silverman M. Trichostatin A-histone deacetylase inhibitor with clinical therapeutic potential-is also a selective and potent inhibitor of gelatinase A expression. Biochem Biophys Res Commun. 2002 Oct 18;298(1):110–5.

63. Harper CV, Finkenstädt B, Woodcock DJ, Friedrichsen S, Semprini S, Ashall L, et al. Dynamic analysis of stochastic transcription cycles. PLoS Biol. 2011;9(4):e1000607.

64. Andrews SS, Brent R. Individual yeast cells signal at different levels but each with good precision. R Soc Open Sci. 2025 Apr 30;12(4):241025.

65. Hansen AS, O’Shea EK. Limits on information transduction through amplitude and frequency regulation of transcription factor activity. eLife. 4:e06559.

66. Pichon X, Lagha M, Mueller F, Bertrand E. A Growing Toolbox to Image Gene Expression in Single Cells: Sensitive Approaches for Demanding Challenges. Mol Cell. 2018 Aug;71(3):468–80.

67. Corrigan AM, Chubb JR. Regulation of Transcriptional Bursting by a Naturally Oscillating Signal. Curr Biol. 2014 Jan 20;24(2):205–11.

68. Fritzsch C, Baumgå rtner S, Kuban M, Steinshorn D, Reid G, Legewie S. Estrogen-dependent control and cell-to-cell variability of transcriptional bursting. Mol Syst Biol. 2018 Feb;14(2). Available from: 10.15252/msb.20177678

69. Rodriguez J, Ren G, Day CR, Zhao K, Chow CC, Larson DR. Intrinsic Dynamics of a Human Gene Reveal the Basis of Expression Heterogeneity. Cell. 2019 Jan;176(1–2):213–226.e18.

70. Zhang Z, Nikolai BC, Gates LA, Jung SY, Siwak EB, He B, et al. Crosstalk between histone modifications indicates that inhibition of arginine methyltransferase CARM1 activity reverses HIV latency. Nucleic Acids Res. 2017 June;45(16):9348–60.

71. Cheng CT, Hsiao JC, Hoffmann A, Tu HL. TNFR1 mediates heterogeneity in single-cell NF-κB activation. iScience. 2024 Apr;27(4):109486.

72. Paszek P, Ryan S, Ashall L, Sillitoe K, Harper CV, Spiller DG, et al. Population robustness arising from cellular heterogeneity. Proc Natl Acad Sci. 2010 June;107(25):11644–9.

73. Pl S, Tp M, E V, Ka J. Histone acetyltransferases regulate HIV-1 enhancer activity in vitro. Genes Dev. 1997 Dec;11(24). Available from: https://pubmed.ncbi.nlm.nih.gov/9407026/

74. Vaid R, Wen J, Mannervik M. Release of promoter–proximal paused Pol II in response to histone deacetylase inhibition. Nucleic Acids Res. 2020 Apr;48(9):4877–90.

75. Jin S, Scotto KW. Transcriptional Regulation of the MDR1 Gene by Histone Acetyltransferase and Deacetylase Is Mediated by NF-Y. Mol Cell Biol. 1998 July;18(7):4377–84.

76. Mohammad IS, He W, Yin L. Understanding of human ATP binding cassette superfamily and novel multidrug resistance modulators to overcome MDR. Biomed Pharmacother. 2018 Apr;100:335–48.

77. Pichon X, Bastide A, Safieddine A, Chouaib R, Samacoits A, Basyuk E, Peter M, Mueller F, Bertrand E. Visualization of single endogenous polysomes reveals the dynamics of translation in live human cells. J Cell Biol. 2016 Sep 12;214(6):769–81.

78. Tsanov N, Samacoits A, Chouaib R, Traboulsi AM, Gostan T, Weber C, et al. smiFISH and FISH-quant - a flexible single RNA detection approach with super-resolution capability. Nucleic Acids Res. 2016 Dec 15;44(22):e165.

79. Stringer C, Wang T, Michaelos M, Pachitariu M. Cellpose: a generalist algorithm for cellular segmentation. Nat Methods. 2020 Dec;18(1):100–6.

80. Imbert A, Ouyang W, Safieddine A, Coleno E, Zimmer C, Bertrand E, et al. FISH-quant v2: a scalable and modular tool for smFISH image analysis. RNA. 2022 Mar;28(6):786–95.

81. Hilfinger A, Paulsson J. Separating intrinsic from extrinsic fluctuations in dynamic biological systems. Proc Natl Acad Sci. 2011 July 19;108(29):12167–72.

82. Fu AQ, Pachter L. Estimating intrinsic and extrinsic noise from single-cell gene expression measurements. Stat Appl Genet Mol Biol. 2016 Dec 1;15(6):447–71.

83. Douaihy M, Topno R, Lagha M, Bertrand E, Radulescu O. BurstDECONV: a signal deconvolution method to uncover mechanisms of transcriptional bursting in live cells. Nucleic Acids Res. 2023 Sept 8;51(16):e88.

84. Radulescu O, Grigoriev D, Seiss M, Douaihy M, Lagha M, Bertrand E. Identifying Markov chain models from time-to-event data: an algebraic approach. Bull Math Biol. 2025;87(1):1–46.

